# TFEB-vacuolar ATPase signaling regulates lysosomal function and microglial activation in tauopathy

**DOI:** 10.1101/2023.02.06.527293

**Authors:** Baiping Wang, Heidi Martini-Stoica, Chuangye Qi, Tzu-Chiao Lu, Shuo Wang, Wen Xiong, Yanyan Qi, Yin Xu, Marco Sardiello, Hongjie Li, Hui Zheng

## Abstract

Transcription factor EB (TFEB) mediates gene expression through binding to the <u>C</u>oordinated <u>L</u>ysosome <u>E</u>xpression <u>A</u>nd <u>R</u>egulation (CLEAR) sequence. TFEB targets include subunits of the vacuolar ATPase (v-ATPase) essential for lysosome acidification. Single nucleus RNA-sequencing (snRNA-seq) of wild-type and PS19 (Tau) transgenic mice identified three unique microglia subclusters in Tau mice that were associated with heightened lysosome and immune pathway genes. To explore the lysosome-immune relationship, we specifically disrupted the TFEB-v-ATPase signaling by creating a knock-in mouse line in which the CLEAR sequence of one of the v-ATPase subunits, *Atp6v1h*, was mutated. We show that the CLEAR mutant exhibited a muted response to TFEB, resulting in impaired lysosomal acidification and activity. Crossing the CLEAR mutant with Tau mice led to higher tau pathology but diminished microglia response. These microglia were enriched in a subcluster low in mTOR and HIF-1 pathways and was locked in a homeostatic state. Our studies demonstrate a physiological function of TFEB-v-ATPase signaling in maintaining lysosomal homoeostasis and a critical role of the lysosome in mounting a microglia and immune response in tauopathy and Alzheimer’s disease.

## Introduction

Lysosomes are intracellular organelles essential for the degradation of protein aggregates and other macromolecules and organelles. Whereas intracellular materials are presented to the lysosome via autophagy, extracellular cargos are taken up through endocytosis or phagocytosis and delivered to the lysosome for clearance. Traditionally regarded as a static organelle for terminal degradation, emerging evidence demonstrates that lysosomes are highly dynamic and tightly regulated^1^. Impaired lysosomal homeostasis has been implicated in aging and age-associated neurodegenerative diseases including Alzheimer’s disease (AD), Parkinson’s disease, and frontotemporal degeneration^2^.

The transcription factor EB (TFEB) plays a central role in lysosome regulation and signaling^3^. It responds to lysosomal pH and content through the LYsosome NUtrient Sensing (LYNUS) machinery composed of v-ATPase, Rag-GTPases and Ragulator, and the recruitment of mTORC1, to undergo cytoplasmic to nucleus trafficking. Inside the nucleus, TFEB promotes the transcription of its target genes through binding to the Coordinated Lysosomal Expression And Regulation (CLEAR) motifs^4, 5^, the network of which consists of genes involved in autophagy, lysosomal biogenesis, lysosomal exocytosis and endocytosis^6^. Thus, TFEB is known as a master regulator of the autophagy and lysosomal pathway. Accordingly, we and others have reported that TFEB overexpression led to the suppression of Aβ and tau pathologies characteristic of AD and other tauopathy diseases in mice^7-12^. While the beneficial effects of exogenous TFEB expression in disease models are abundantly documented, the role of endogenous TFEB in AD pathogenesis is less well-defined. Further, whether these effects are solely mediated through lysosomal clearance remains unclear.

A key determinant of the lysosomal functionality is its acidic pH controlled by the v-ATPase^13^. Reduced v-ATPase activity and defective lysosomal acidification have been implicated as early events in AD progression^2^. In addition to promoting the expression of a broad range of lysosomal enzymes, TFEB targets also include subunits of the v-ATPase^6^. Of interest, in *Drosophila*, TFEB homologue MITF exclusively regulates the v-ATPase subunits^14, 15^, indicating evolutionary conservation of the TFEB-v-ATPase regulatory pathway. We found that the v-ATPase and lysosomal pathway as well as the immune pathway genes were prominently upregulated in the PS19 tau transgenic (herein referred to as Tau) mouse brains. Through manipulating the endogenous TFEB-v-ATPase signaling, executed by mutagenesis of the CLEAR sequence in the promoter of one of the v-ATPase subunits, *Atp6v1h*, we demonstrate that specific disruption of the TFEB-dependent *Atp6v1h* transcriptional regulation leads to impaired v-ATPase activity and lysosomal function under physiological conditions. Intriguingly, microglia with the disrupted TFEB-v-ATPase signaling fail to be activated in Tau mice, revealing an essential role of the lysosome in initiating microglia and immune pathway activation.

## Results

### Upregulated TFEB and lysosomal pathway in tauopathy

Our previous work revealed that TFEB and several of its lysosomal target genes were significantly increased in human tauopathy brain samples and in Tau mice^12^. To investigate this phenomenon further, we conducted hippocampal bulk RNA-seq in wild-type (WT) and Tau mice either before (4 months) or after (9 months) the development of tangle-like pathologies (Supplementary Table 1). We found only a few differentially expressed genes (DEGs) between WT and Tau mice at 4 months of age. In contrast, we identified 825 significantly upregulated genes and 89 significantly downregulated genes (cutoff of FDR < 0.05 and Fold Change >1.5) in 9-month-old Tau mice compared to WT (Extended Data Fig. 1a-d). These were validated by quantitative PCR (qPCR) analysis (Extended Data Fig. 1e,f). These results indicate that the DEGs identified in 9-month-old Tau samples were induced by tau pathology rather than transgene overexpression. Gene set enrichment analysis (GSEA) revealed highly significant enrichment of both the lysosome and inflammatory response pathway genes in Tau mice (Fig. 1a,b).

**Figure 1.**
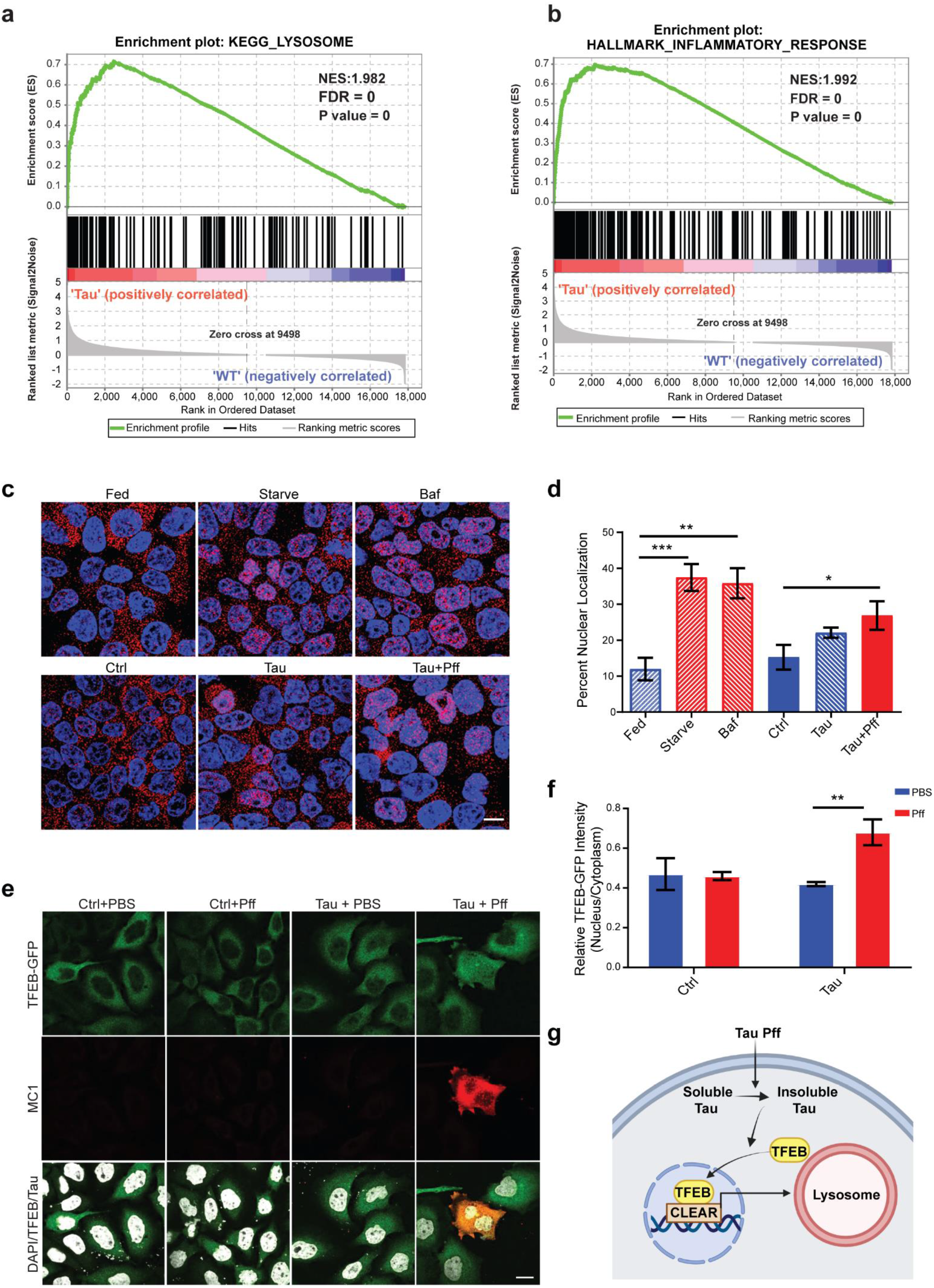
Elevated TFEB and lysosomal pathway in PS19 (Tau) mice and by insoluble tau. **a**,**b**. Gene set enrichment analysis (GSEA) of lysosome (a) and inflammatory (b) pathways in Tau mice (n=4) compared with WT mice (n=5) at 9 months. **c**. Representative fluorescence images of HEK293 cells transfected with empty vector (Ctrl) or Tau-P301L expression vector (Tau) or Tau expressing cells treated with Pff (Tau + Pff) and immune-stained with an anti-TFEB antibody for endogenous human TFEB (red) and DAPI (blue). Cells under normal growth (Fed) condition were used as a negative control whereas cells grown in serum-free medium (Starve) or treated with 200 nM Bafilomycin (Baf) were used as positive controls. **d**. Quantification of percent TFEB nuclear localization showing significantly higher nuclear TFEB in Tau+Pff group. N=13 images/condition. **e**. Representative fluorescence images of HEK293 cells co-transfected with TFEB-GFP plus empty vector (Ctrl) or Tau-P301L construct and treated with PBS or Pff, followed by staining with the MC1 antibody, showing prominent nuclear TFEB in MC1 positive cells. **f**. Quantification of TFEB-GFP nuclear/cytoplasmic ratio, showing significantly higher nuclear TFEB in Tau+Pff cells. N=10-15 images/condition. **g**. A working model whereby Tau Pff converts cellular Tau from soluble to insoluble form, which in turn induces TFEB nuclear translocation and upregulation of lysosomal gene expression. Scale bar: 10 µm. Data are presented as average ± SEM. Two-tailed Student’s *t-*test. *p<0.05, **p<0.01, ***p<0.001. The experiments were repeated 3 times with each in triplicates. See also Extended Data Fig. 1.

To directly test whether tau pathology induces TFEB activation and lysosomal gene expression, we first examined endogenous TFEB localization in HEK293 cells in response to tau. When the cells were fed with normal serum-containing medium (Fed), TFEB was predominantly expressed in the cytoplasm, but translocated to the nucleus when the cells were serum starved (Starve) or treated with Bafilomycin (Baf), a v-ATPase inhibitor that induces lysosomal stress (Fig. 1c,d). We transfected either the empty vector (Ctrl) or the P301L mutant tau (Tau) to HEK 293 cells and a portion of the Tau cells were seeded with tau pre-formed fibrils (Pff), which converts soluble tau to insoluble aggregates^16^. Immunostaining for endogenous TFEB showed that compared to vector-transfected controls, the Tau expressing cells showed a trend of higher percentage of nuclear TFEB, and this became significant when the Tau cells were seeded with Pff (Tau+Pff) (Fig. 1c,d), indicating tau aggregation induces endogenous TFEB nuclear translocation. This was further validated by co-transfecting TFEB-GFP with either the empty vector (Ctrl) or the P301L tau (Tau) to HEK293 cells, followed by treating the cells with either the vehicle (PBS) or tau Pff. Immunostaining with the tau confirmation antibody MC1 (Fig. 1e) followed by quantification (Fig. 1f) showed that MC1-positive cells in Tau+Pff group displayed significantly higher nuclear TFEB compared to other conditions. These results support a model whereby seeding-induced insoluble tau triggers TFEB nuclear translocation and downstream lysosomal gene expression (Fig. 1g).

### snRNA-seq revealed drastically altered microglial profiles in Tau mice

Having established upregulated lysosomal and immune pathways in bulk brains of Tau mice, we next sought to understand the cell types contributing to the changes by conducting single-nucleus RNA-sequencing (snRNA-seq) of the hippocampus collected from 9-month-old WT and Tau mice. Nuclei isolated by fluorescence activated cell sorting (FACS) were profiled using the droplet-based 10x Genomics platform. After stringent quality control including doublet removal, batch effect correction and normalization (Extended Data Fig. 2a), we obtained a total of 55,254 high-quality single cell transcriptomes (Supplementary Table 2), which were annotated into 8 major cell types based on the expression of well-known cell-type-specific markers (Fig. 2a-c). Cell type composition analysis between WT and Tau mice revealed that certain neuronal populations, particularly granule cell cluster, were strongly reduced in Tau mice, indicating neurodegeneration (Fig. 2b). In contrast, the microglia population was greatly expanded in Tau mice. Further analysis identified 915 DEGs in the microglia of Tau mice compared with WT mice, of which 600 were significantly upregulated genes with a cutoff of FDR < 0.05 and log_2_ Fold Change >0.25. We found that signatures of disease-associated microglia (DAM) and microglia of neurodegeneration type (MGnD)^17, 18^ were among the top upregulated DEGs (Fig. 2d). Gene Ontology (GO) pathway analysis of the upregulated genes revealed immune and lysosome pathways as top enriched pathways in Tau microglia (Fig. 2e).

**Figure 2.**
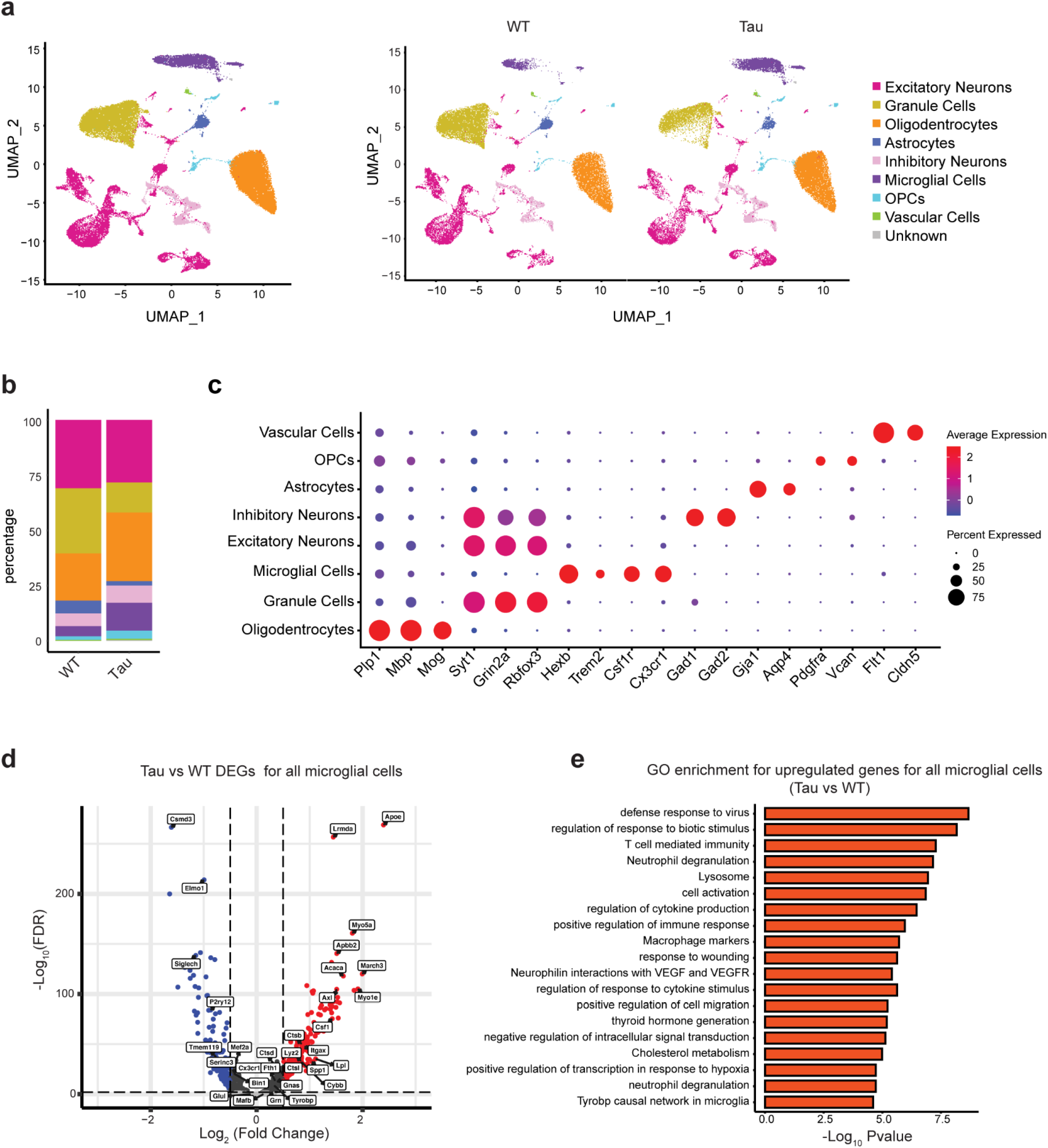
snRNA-seq revealed lysosome and immune pathway upregulation in microglia of Tau mice. **a**. UMAP representation of snRNA-seq analysis of 55,254 cells from hippocampus of WT and Tau mice (left panel) and across genotypes (right panel). Cell type annotation was based on the expression of markers shown in panel (c). **b**. Stacked barplot showing the cell type compositions comparing Tau with WT. **c**. Average scaled expression levels of selected signature genes for different cell types. **d**. Volcano plot showing differentially expressed genes (DEGs) for all microglia in Tau versus WT mice. Up-regulated genes are highlighted in red, Down-regulated genes are highlighted in blue. **e**. Gene ontology enrichment analysis of biological processes for up-regulated expressed genes in microglia of Tau versus WT mice. See also Extended Data Fig. 2.

Further analysis of the microglia population identified seven subclusters (0-6, Fig. 3a). Subcluster 0 was mostly highly represented in WT microglia (Fig. 3b), which has the characteristics of homeostatic microglia with higher expression of *P2ry12, Ccr5, Siglech* (Fig. 3c and Extended Data Fig. 2b). This subpopulation was drastically reduced while subcluster 1 was greatly expanded in Tau microglia. In addition, the Tau microglia gained two unique populations: subclusters 2 and 3 (Fig. 3a,b). Subclusters 1, 2 were enriched for DAM signatures, such as *Apoe, Axl, Csf1*, while subcluster 3 resembled interferon (IFN)-responsive microglia with highest expression of *Oasl2, Ifi204 and Ifi207* (Fig. 3c and Extended Data Fig. 2b). Of note, we did not detect changes of *Trem2* in Tau microglia (Fig. 3e). Comparisons of upregulated DEGs between subclusters 1, 2 and 3 with 0 revealed that most of the subcluster 1 DEGs were included in subclusters 2 and 3 while subclusters 2 and 3 displayed distinct DEGs (Fig. 3d). Thus, subcluster 1 presents as an intermediate state while subclusters 2 and 3 acquired distinct features. GO term enrichment analysis of upregulated genes between subcluster 2 and 0 revealed that lysosome and inflammatory response pathways were strongly over-represented (Fig. 3e,f), while comparison between subcluster 3 and 0 identified anti-virus and interferon responsive pathways as top enriched pathways, confirming subcluster 3 as interferon responsive microglia (Fig. 3g,h). Other smaller clusters were not further characterized.

**Figure 3.**
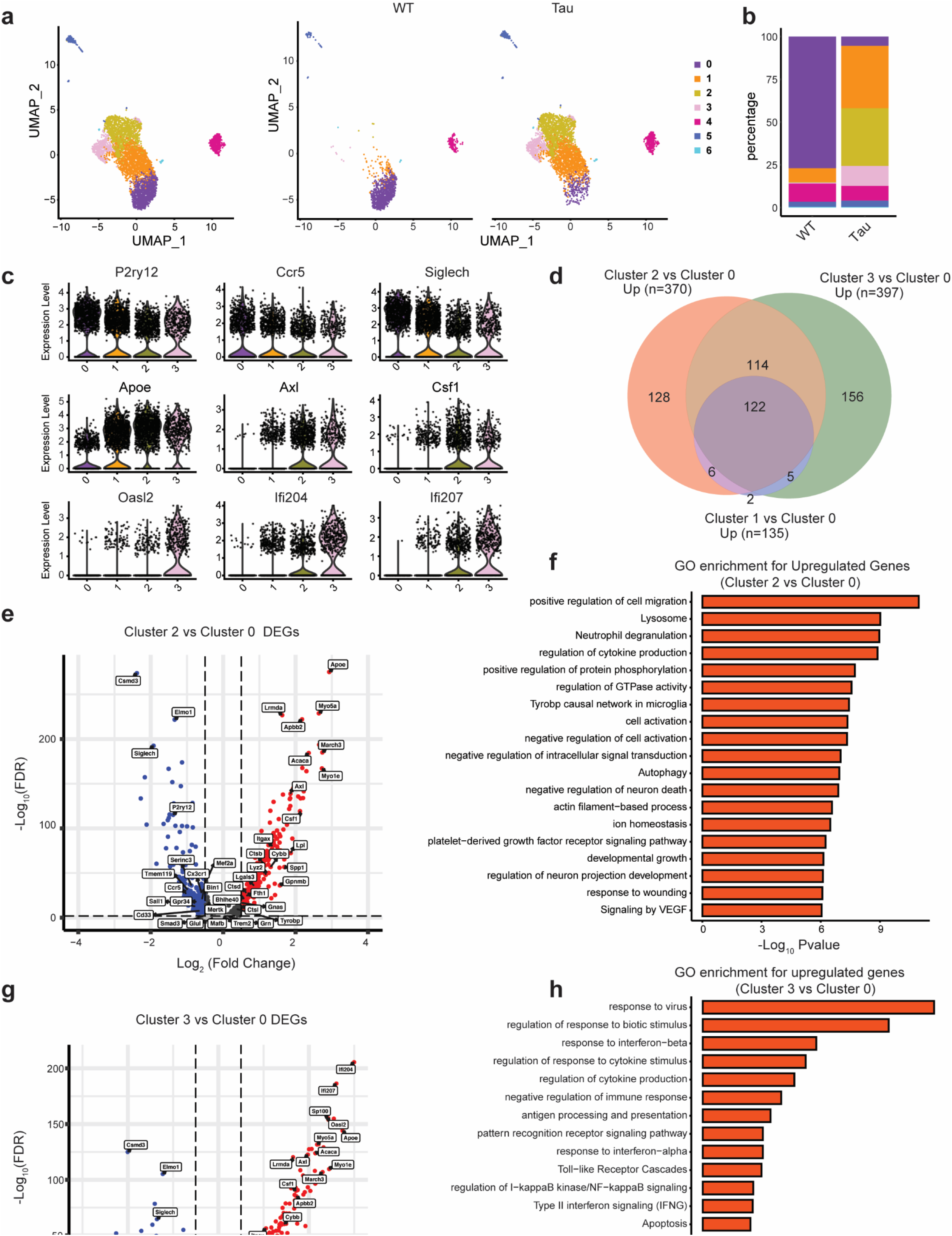
Drastic shift of microglia subclusters in Tau mice. **a**. UMAP representation of microglia subclusters (left panel) and the subclusters across genotypes (right panel). **b**. Stacked barplot showing the subcluster compositions of microglia comparing Tau with WT. **c**. Violin plot showing the expression level of homeostatic microglia genes (P2ry12, Ccr5 and Siglech), disease-associated-microglia genes (Apoe, Axl and Csf1) and IFN responsive-microglia genes (Oasl2, Ifi204 and Ifi207) in subclusters 0, 1, 2 and 3. **d**. Venn diagram summarizing the numbers of up-regulated genes in subclusters 1, 2 and 3 versus 0. **e**. Volcano plot showing differentially expressed genes (DEGs) between subcluster 2 and subcluster 0. Up-regulated genes are highlighted in red, Down-regulated genes are highlighted in blue. **f**. Gene ontology enrichment analysis of biological processes for up-regulated expressed genes in subcluster 2 versus 0. **g**. Volcano plot showing DEGs for microglia subcluster 3 versus subcluster 0. Up-regulated genes are highlighted in red, Down-regulated genes are in blue. **h**. Gene ontology enrichment analysis of biological processes for up-regulated expressed genes in microglia of subcluster 3 versus 0.

### In vivo modeling of TFEB-v-ATPase signaling through CLEAR mutagenesis

The fact that the lysosomal pathway genes were prominently enriched in both the total microglia and subcluster 2 of Tau mice prompted us to seek further understanding of its functional implications and its relationship with the immune pathway. Although this could be achieved by TFEB manipulation, the many other non-lysosomal genes TFEB also targets, in particular the immune pathway genes^19^, makes it difficult to delineate the lysosome-specific effect. We thus zoomed in on the TFEB lysosomal-specific target, the v-ATPase, given its key role in regulating lysosomal pH and activity. We found that most of the v-ATPase subunit genes were upregulated in Tau microglia (Fig. 4a), and hypothesized that disruption of TFEB-v-ATPase transcriptional regulation through mutagenizing the TFEB-binding CLEAR motif may lead to reduced v-ATPase activity and impaired lysosomal function. We chose the *Atp6v1h* subunit gene as it contains two strong tandem repeat CLEAR sequences within its promoter region (Fig. 4b). Chromatin immunoprecipitation (ChIP) of N2a cells transfected with GFP-FLAG or TFEB-FLAG using an anti-FLAG antibody followed by qPCR confirmed TFEB binding to the CLEAR sequence of the *Atp6v1h* promoter (Fig. 4c). *Mcoln1* was used as a positive control while Chr 1, 2, and 3 representing gene deserts of respective chromosomes lacking CLEAR sequences were used as negative controls. To validate that TFEB-CLEAR interaction promotes transcriptional activation, we cloned either the wild-type (WT) or the CLEAR mutant (CL) *Atp6v1h* promoter fragments to the firefly luciferase reporter, using the CLEAR-lacking AQP1-luciferase as a negative control, and co-transfect the constructs with either empty vector (CMV) or a TFEB expression vector. The luciferase assay showed that, compared with CMV controls, the WT *Atp6v1h* promoter responded to TFEB as expected, the TFEB response was blunted in the CL mutant as with the AQP1 control (Fig. 4d). Consistent with our earlier results that insoluble tau induces TFEB activation (Fig. 1c-f), addition of tau Pff to tau-expressing cells (Tau+Pff) also induced the luciferase activity driven by the WT *Atp6v1h* promoter (Fig. 4e,f), and this effect was blocked when the CL mutant promoter was used (Fig. 4f). These data combined demonstrate that TFEB binds to the CLEAR sequence of the *Atp6v1h* promoter and activates its gene expression. Insoluble tau induces TFEB nuclear translocation and enhances *Atp6v1h* transcription in a CLEAR dependent manner.

**Figure 4.**
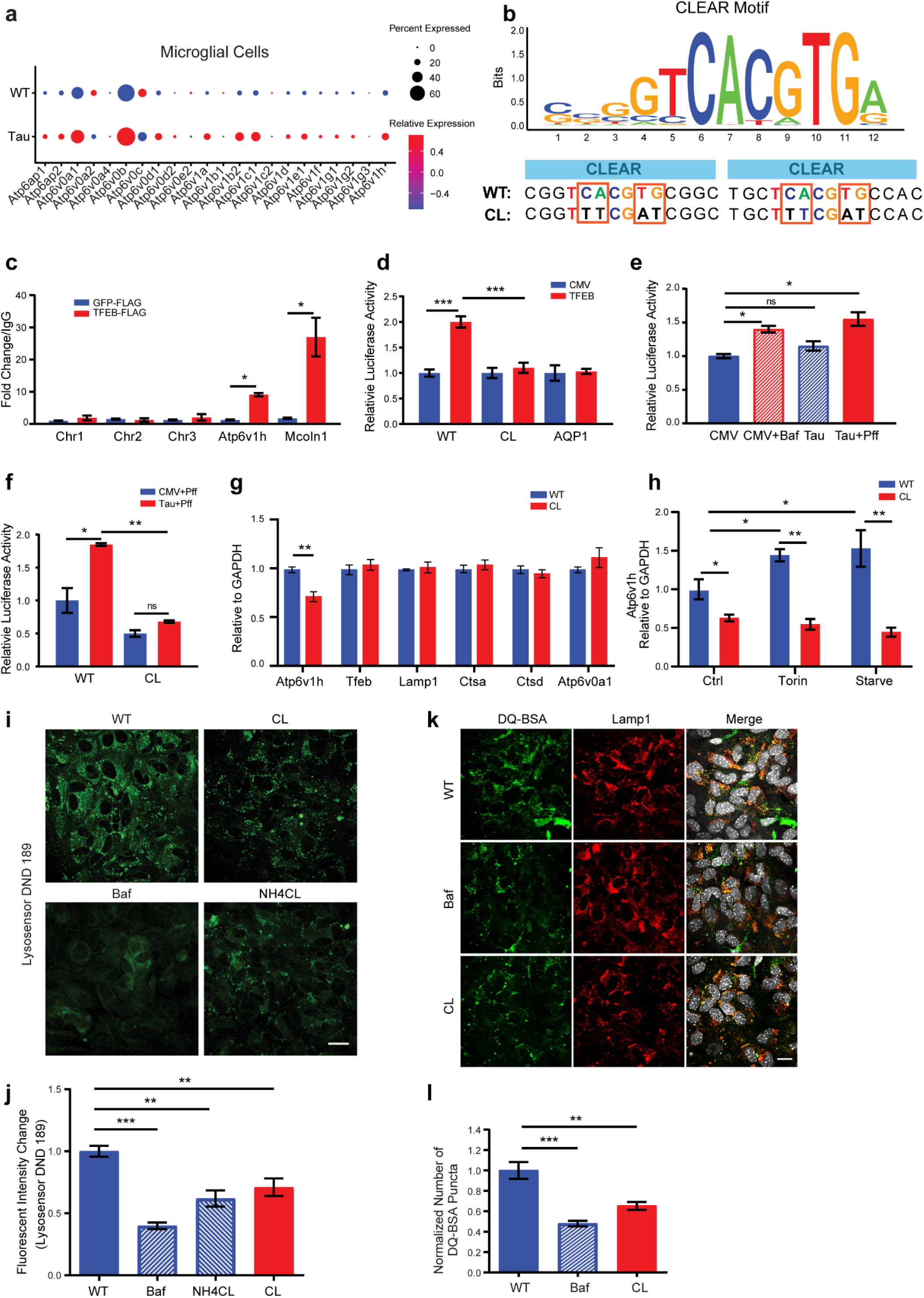
Generation of an in vivo model of lysosomal dysfunction through CLEAR mutagenesis. **a**. Dotplot showing the relative gene expression levels of the v-ATPase subunits in microglia of WT and Tau mice. **b**. The consensus TFEB binding CLEAR motif (upper panel). **WT**: two canonical CLEAR sequences in the promoter of *Atp6v1h*. **CL**: Mutated CLEAR motif with the altered base pairs highlighted. **c**. TFEB binds the *Atp6v1h* promoter region predicted to contain CLEAR sequence. Chromatin immunoprecipitation and qPCR analysis (CHIP-PCR) of N2a cells transfected with TFEB-FLAG or GFP-FLAG with an anti-FLAG antibody. *Mcoln1*, a known TFEB target with multiple CLEAR sequences in the promoter, was used as a positive control. Chr 1, 2, and 3 represent gene deserts in those respective chromosomes containing no CLEAR sequences were used as negative controls. **d**. CLEAR sequence mutagenesis in Atp6v1h ablates TFEB transcriptional activity. Luciferase assay in HEK293 cells co-transfected with TFEB, WT or CL mutant Atp6v1h promoter driven firefly luciferase construct and the Renilla construct. CMV vector was used as a negative control. Firefly luciferase activities were normalized to Renilla. AQP1 is a promoter construct that does not contain a CLEAR sequence and not regulated by TFEB. **e**. Luciferase assay demonstrating that insoluble tau promotes Atp6v1h transcription. Cells were co-transfected with empty vector (CMV) or Tau-P301L vector (Tau) with the wild type Atp6v1h promoter luciferase and Renilla vectors. Tau+Pff: Pff was added for seeding insoluble tau. Cells treated with 200 nM Bafilomycin (Baf) were used as a positive control. **f**. The same luciferase assay showing that Tau+Pff failed to activate the luciferase activity when the CL protomer was used demonstrating that insoluble tau enhances Atp6v1h promoter activity in a CLEAR sequence dependent manner. **g**. CL mutant mice exhibit a specific reduction in Atp6v1h transcript without affecting other TFEB lysosomal targets. qPCR analysis of forebrain RNA extract from 1-month-old mice homozygous for CLEAR mutant (CL) or WT control. N=4/group. **h**. qPCR analysis of Atp6v1h transcripts in WT and CL primary glial cultures under basal (Ctrl), Torin treated or starvation (Starve) conditions, showing reduced Atp6v1h expression under Ctrl conditions and were unresponsive to Torin or starvation treatments. **i**. Representative images of LysoSensor Green DND-189 fluorescence in WT and CL primary glial cultures. Bafilomycin (Baf) and NH_4_Cl treated WT cultures were used as controls. **j**. Quantification of (i) showing reduced lysosomal acidification in CL cultures. **k**. Representative images of DQ-BSA fluorescence co-stained with Lamp1 in WT and CL primary glial cultures. Bafilomycin (Baf) treated WT cultures were used as a control. **l**. Quantification of (k) showing reduced lysosomal degradation capacity in CL cultures. Data are presented as average ± SEM. *p<0.05, **p<0.01, ***p<0.001 by 1-way ANOVA with Sidak’s correction. Each in vitro experiment was repeated 3 times with each in triplicates.

To test the functional role of the TFEB-*Atp6v1h* signaling in lysosomal regulation and tauopathy in vivo, we introduced the same CLEAR mutation into the endogenous mouse *Atp6v1h* promoter via CRISPR/Cas9 technology. Mice homozygous for the CLEAR mutation (CL) showed a 25-30% reduction of the *Atp6v1h* transcript in whole brain extracts while neither TFEB nor other TFEB targets were affected (Fig. 4g). To directly validate that the CLEAR mutant obliterates TFEB’s transcriptional regulation of *Atp6v1h*, we prepared primary mixed glia cultures from WT and CL homozygotes and treated them with Torin or starvation that are known to induce TFEB nuclear localization and activation of its target genes^20-22^. In line with the bulk brain PCR, *Atp6v1h* transcripts were reduced by approximately 40% in CL homozygote cultures (Fig. 4h, Ctrl). In WT cultures, TFEB activation in both Torin and starvation treated conditions enhanced *Atp6v1h* transcription, while in CL cultures, the induction of *Atp6v1h* transcripts through these treatments was abolished (Fig. 4h). Collectively, these results reveal a physiological regulation of *Atp6v1h* transcription by TFEB through the CLEAR sequence in vivo and in vitro and the specific disruption of TFEB-*Atp6v1h* signaling without affecting other TFEB targets in the CL mutant mice. Next, we sought to determine whether the disrupted TFEB-*Atp6v1h* regulation leads altered v-ATPase activity and lysosomal function. We first measured the lysosomal acidity in primary mixed glia cultures from WT and CL mice using Lysosensor Green DND-189, a pH sensitive dye that exhibits increased fluorescence in acidic organelles. For positive controls, we treated WT cultures with Bafilomycin (Baf) or NH_4_CL which are known to elevate lysosomal pH. Compared with the WT control, the CL cultures showed significantly reduced Lysosensor fluorescence (Fig. 4I,j), indicating a defect in lysosomal acidification. To determine the functional consequences of reduced acidification in CL cultures, we utilized DQ-bovine serum albumin (DQ-BSA), which becomes fluorescent upon degradation, to assay the overall lysosomal hydrolase activity. Using Baf-treatment as a positive control, we showed that the intensity of DQ-BSA fluorescence was significantly decreased in CL cells compared with the WT (Fig. 4k,l), indicating reduced lysosomal degradative capacity. Thus, disruption of the TFEB-*Atp6v1h* signaling in the CLEAR mutant ablates v-ATPase activity, leading to defective lysosomal acidification and degradation.

### Increased tau pathology but reduced glial activation in Tau mice crossed with the CL mice

Having successfully created an in vivo model of lysosomal dysfunction, we next interrogated the role of TFEB-v-ATPase regulation in tauopathy by crossing the CL mice with the Tau mice followed by analysis at 9 month-of-age. Western blotting of the forebrain hemispheres of Tau and Tau;CL mice showed a significant increase of phospho-tau species identified by PHF1 and CP13 antibodies in Tau;CL mice compared with Tau mice (Fig. 5a,b). These results were further confirmed by immunostaining of the brain slices with the AT8 antibody showing Tau;CL mice exhibited increased AT8 positive phospho-tau compared to Tau mice (Fig. 5c,d). Surprisingly, co-immunostaining for GFAP and Iba1 showed that, despite increased phospho-tau pathology in Tau;CL mice, levels of astrogliosis, measured by GFAP fluorescence intensity (Fig. 5e) and microgliosis, quantified by Iba1 fluorescence and microglia number (Fig. 5f,g), were significantly reduced. Further examination of microglia morphologies by 3D reconstruction and IMARIS analysis revealed that both the surface area and volume were reduced in Tau;CL compared with Tau mice (Fig. 5h-j), providing additional support for their dampened response to tauopathy.

**Figure 5.**
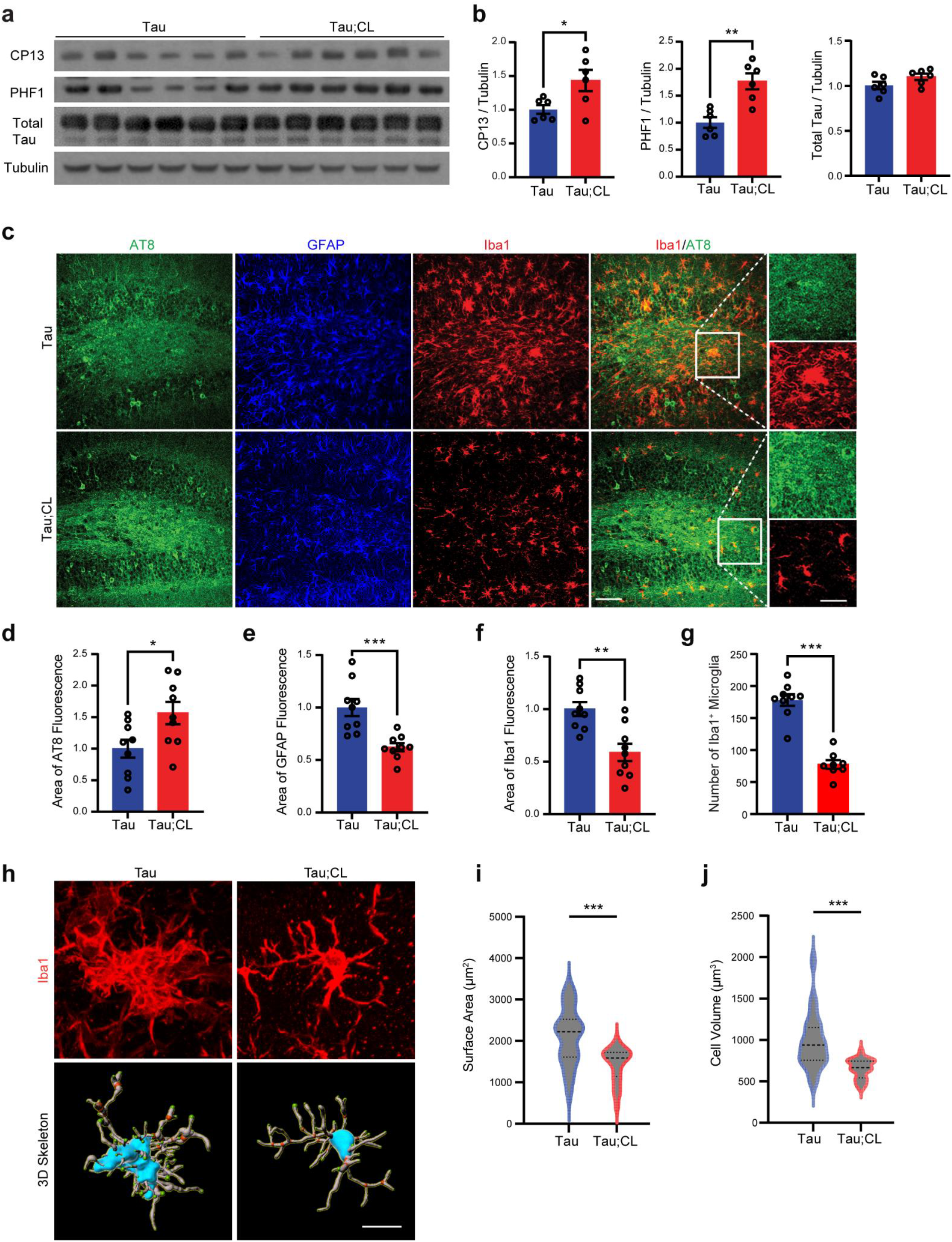
Increased phospho-tau and decreased gliosis in Tau mice crossed to CL background. **a**,**b**. Western blot (a) with quantification (b) of total and phospho-tau species recognized by PHF1 and CP13 antibodies from forebrain lysates of 9-month-old Tau mice or Tau mice homozygous for the CL mutation (Tau; CL). N=6/group. **c-g**. Representative fluorescent confocal images of AT8, GFAP and Iba1 immunostaining (c) with quantification of AT8 (d), GFAP(e) and Iba1 (f) fluorescence intensities and Iba1 positive cells (g) in the dentate gyrus samples of 9-month-old Tau and Tau;CL) mice. Scale bar 100 µm and 50 µm in brackets. N=8/group. **h**. Representative Iba1 staining and 3D skeletonization of microglia in the hippocampus of Tau and Tau;CL mice. Scale bar:10 um. **i**,**j**. Quantification of microglia surface area (i) and volume (j) per cell using the IMARIS software. N=6/group. Data are presented as average ± SEM. Two-tailed Student’s *t-*test. *p<0.05, **p<0.01, ***p<0.001. See also Extended Data Fig. 3.

Since the CL mice had ∼30% reduction in *Atp6v1h* levels, we wondered whether the phenotypes observed in Tau;CL mice were attributed by decreased *Atp6v1h* expression or due to disrupted TFEB-*Atp6v1h* signaling regulation. To address this question, we created an *Atp6v1h* germline heterozygous knockout (VKO) allele and crossed the mice with the Tau mice (Extended Data Fig. 3). qPCR analysis showed a ∼50% reduction of *Atp6v1h* transcript in the brains of VKO and Tau19;VKO mice (Extended Data Fig. 3a). Immunostaining using AT8 and anti-GFAP and -Iba1 antibodies revealed comparable levels of phospho-tau intensity and gliosis between the Tau and Tau;VKO mice (Extended Data Fig. 3b-e). This was further validated by Western blotting of total and phospho-tau levels (Extended Data Fig. 3f,g) and qPCR analysis of TNF*α* and IL1β expression (Extended Data Fig. 3h,i). These results highlight the importance of TFEB-v-ATPase signaling regulation rather than the basal level of *Atp6v1h* expression in microglia reactivity and tauopathy.

To assess whether the changes in the Tau;CL microglia were due to the intrinsic defects in the CL mice, we performed 3D reconstruction of WT and CL microglia (Fig. 6a). We found that the CL microglia had reduced total processes, surface area, cell volume and terminal and branch points compared to WT controls (Fig. 6b). To further evaluate the functional role of these changes in immune activation, we performed *i*.*p* injection of LPS to WT and CL mice and measured the expressions of proinflammatory cytokine in hippocampal samples. The CL mutant mice showed significantly less induction of TNF*α*, IL1β, IL6 and Irf7 compared to WT controls (Fig. 6c), indicating compromised immune responses in CL mutant. This was also the case when primary microglia cultures were challenged with LPS (Fig. 6d).

**Figure 6.**
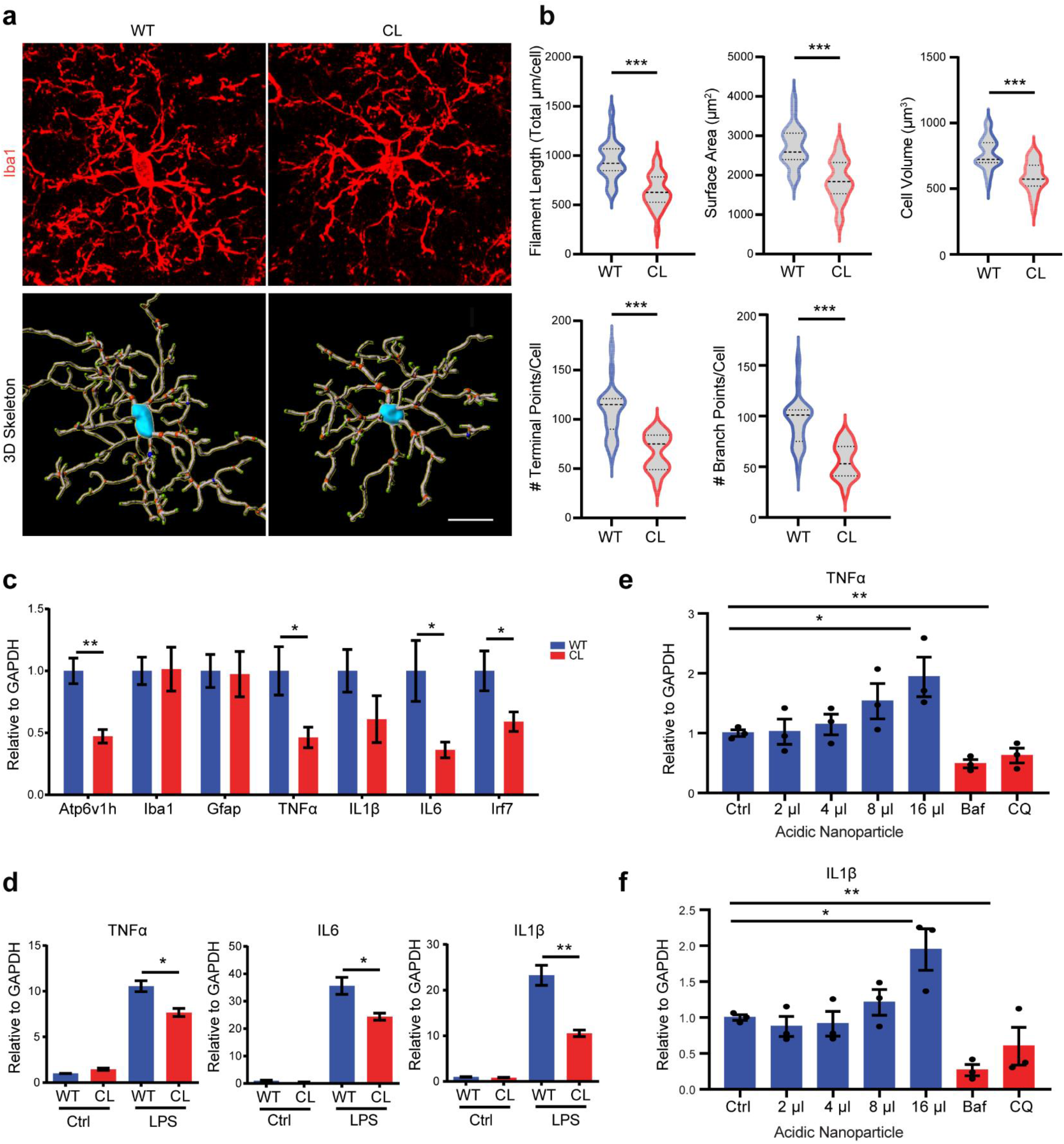
Disruption of TFEB-v-ATPase lysosomal signaling leads to Impaired microglia morphology and activation. **a**. Representative Iba1 staining and 3D skeletonization of microglia in the hippocampus of WT and CL mice. Red dots marked branching points of microglia processes; green dots marked terminal points. Scale bar:10 um. **b**. Quantification of microglia filament length, surface area, volume, terminal points and branch points using the IMARIS software. N=4/group. **c**. qPCR analysis of proinflammatory cytokine expressions in hippocampus tissues of 4-month-old WT and CL mice injected with LPS. N=4/genotypes. **d**. qPCR analysis of proinflammatory cytokine expressions in primary microglia cultures generated from WT and CL pups at basal conditions (Ctrl) and with LPS stimulation. **e**,**f**. qPCR analysis of TNF*α* (e) or IL1β (f) levels of primary microglia cultures treated with LPS together with increasing doses of acidic nanoparticles to increase lysosomal acidity, or Bafilomycin (Baf) or Chloroquine (CQ) to reduce lysosomal acidity, showing that modulation of lysosomal acidity directly leads to altered proinflammatory cytokine expressions. Data are presented as average ± SEM. Two-tailed Student’s *t-*test. *p<0.05, **p<0.01, ***p<0.001. Each in vitro experiment was repeated 3 times with each in triplicates. See also Extended Data Fig. 4.

To test whether changing lysosomal acidity can directly modulate immune response, we treated primary mixed glia cultures with acidic nanoparticles to acidify the lysosome^23^. Immunostaining showed that the nanoparticles were delivered to Lamp1 positive lysosomes (Extended Data Fig. 4). Co-treatment of the acidic nanoparticles with LPS showed dose-dependent increases of TNFα and IL1β expressions (Fig. 6e,f). In contrary, co-treatment of LPS with Bafilomycin (Baf) or Chloroquine (CQ), both of which are known to increase lysosomal pH, led to greatly diminished TNFα and IL1β expression (Fig. 6e,f). These results demonstrate a direct regulation of the immune response by lysosomal pH. Overall, we have established that proper lysosomal acidification mediated by TFEB-v-ATPase signaling is essential for immune activation.

### snRNA-seq identified a unique mTOR and HIF1 low microglia subcluster with dysregulated TFEB-v-ATPase signaling that is locked in the homeostatic state

To understand the molecular mechanisms by which TFEB-v-ATPase signaling regulates microglia activity, we carried out snRNA-seq analysis of hippocampus obtained from 9-month-old WT, CL, Tau and Tau;CL littermates (Fig. 7a, Extended Data Fig. 5a,b). Cell composition analysis showed that, consistent with the Iba1 immunostaining, the expanded microglia population in Tau mice was substantially reduced in Tau;CL (Fig. 7b). Further clustering of microglia identified 10 distinct subclusters (Fig. 7c). Based on the expression of marker genes described in Fig. 3, we were able to further divide the homeostatic subcluster 0 into two subpopulations, 0a and 0b, which together with the previously identified subclusters 1 (transitional), 2 (DAM/MGnD-like), and 3 (IFN responsive), consist of the major microglia subclusters and were analyzed further. Composition analysis of each subcluster across genotypes revealed that subcluster 0a was reduced whereas 0b was expanded in CL mice compared to WT and both were greatly diminished in Tau samples (Fig. 7d,e). Strikingly, subcluster 0b was largely preserved in Tau;CL mice, indicating that this subcluster was unable to be converted to activated states. This effect was observed in both male and female mice, despite slight gender differences in microglia profiles observed in Tau mice (Extended Data Fig. 5c). Analysis of DEGs between WT and CL microglia showed that the majority of genes were down-regulated (Extended Data Fig. 6), suggesting that the expanded subcluster 0b was associated with suppressed gene expression profiles. Further analysis of subcluster 0b with 0a revealed that the downregulated genes include the lysosome (Fig. 7g), mTOR (Fig. 7h) and Hypoxia Inducible Factor-1 (HIF-1) (Fig. 7i) signaling pathways. In contrast, these pathway genes were prominently upregulated in both subclusters 2 and 3, and to a less degree in subcluster 1, compared with subcluster 0a (Fig. 7g-i). These results indicate that microglia in subcluster 0b were refractory to initiate the activation process in tauopathy conditions, due to lower lysosomal activity and possibly associated mTOR and HIF-1 signaling pathways (Fig. 7j).

**Figure 7.**
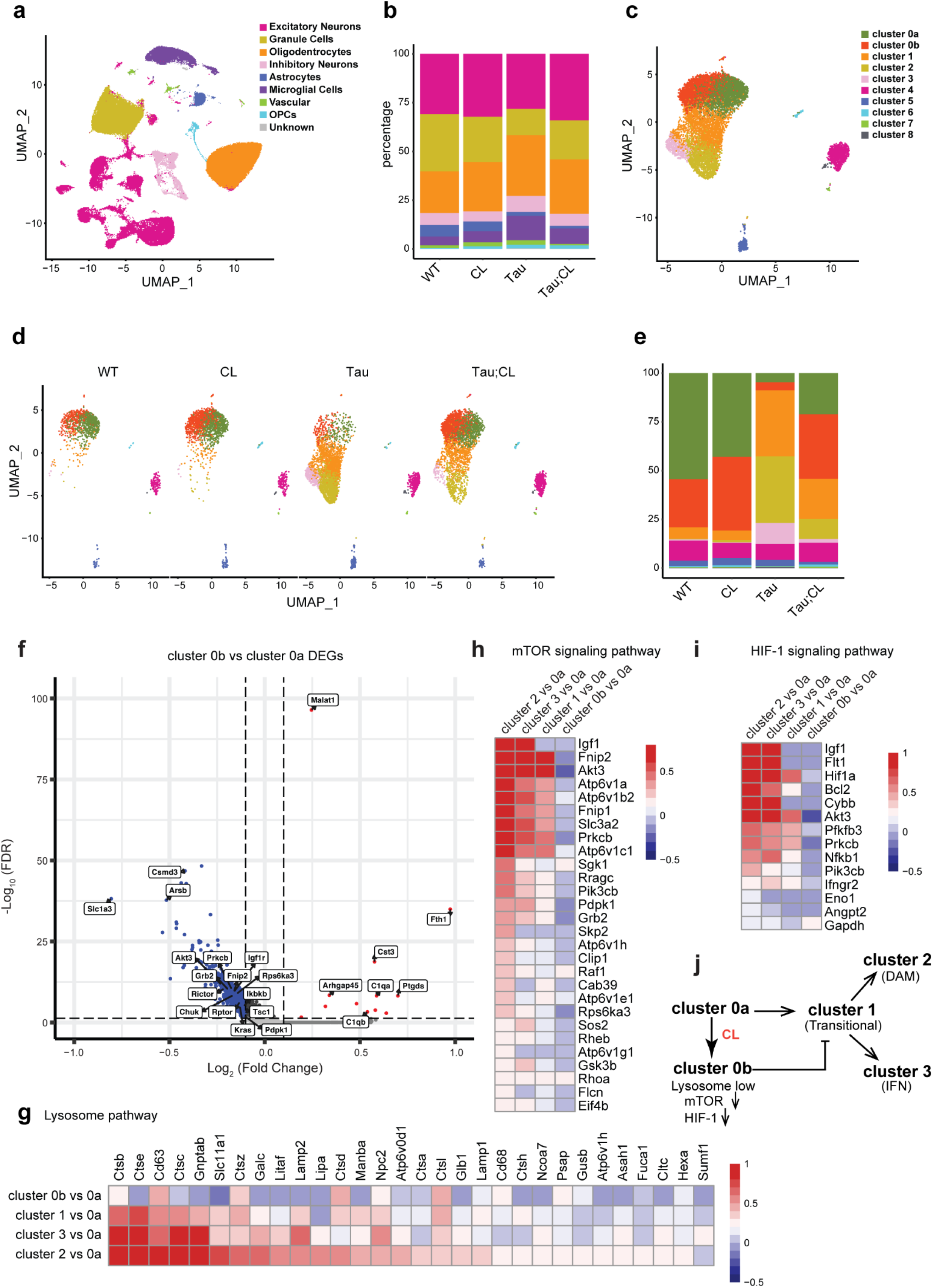
snRNA-seq analysis identified a distinct homeostatic microglia subcluster regulated by TFEB-vATPase. **a**. UMAP representation of snRNA-seq analysis of 137,734 cells from hippocampus of WT, CL, Tau and Tau;CL mice. **b**. Stacked barplot showing cell compositions across different genotypes. **c**. UMAP representation of re-clustered microglia cells, with further separation of subcluster 0 to 0a and 0b. **d**. UMAP representation of re-clustered microglia cells across genotypes. **e**. Stacked barplot showing subcluster compositions of microglia across different genotypes. **f**. Volcano plot showing differentially expressed genes (DEGs) for cluster 0b versus cluster 0a. Up-regulated genes are highlighted in red, Down-regulated genes are highlighted in blue. Significantly downregulated DEGs in mTOR pathway are labeled. **g-i**. The heatmaps comparing the levels of lysosome (g), mTOR (h) and HIF-1 (i) signaling pathways related genes (log_2_ fold change) in different microglia subclusters. **j**. A model illustrating microglia subcluster relationships. Microglia in subclusters 0b with lower lysosomal, mTOR and HIF-1 activities are refractory to transition toward activated microglia subcluster 2 and 3 upon activation; Loss of TFEB-vATPase regulation (CL) drives the expansion of homeostatic subcluster 0b at the expanse of subcluster 0a. See also Extended Data Figs. 5 and 6.

Consistent with this notion, comparison between Tau and Tau;CL microglia showed that the preservation of subcluster 0b in Tau;CL was correlated with drastically reduced subclusters 1, 2 and 3 (Fig. 7,d,e). Accordingly, the upregulated DEGs identified in Tau microglia, including lysosome, immune and lipid metabolic pathway genes were markedly reduced in Tau;CL (Fig. 8a,b and Extended Data Fig. 7). In agreement with the subcluster analysis, the mTOR and HIF-1 signaling pathways were upregulated in Tau but downregulated in CL and Tau;CL microglia (Fig. 8c,d). Given the prominent role of the mTOR and its downstream HIF-1 signaling pathways in mediating cellular metabolism and immune cell activation, we performed immunostaining of Hif1a, a subunit of HIF-1, on brain slices from 9-month-old WT, CL, Tau, and Tau;CL mice. Our results demonstrate upregulated Hif1a expression in Tau microglia, which was significantly reduced in Tau;CL microglia (Fig. 8e,f).

**Figure 8.**
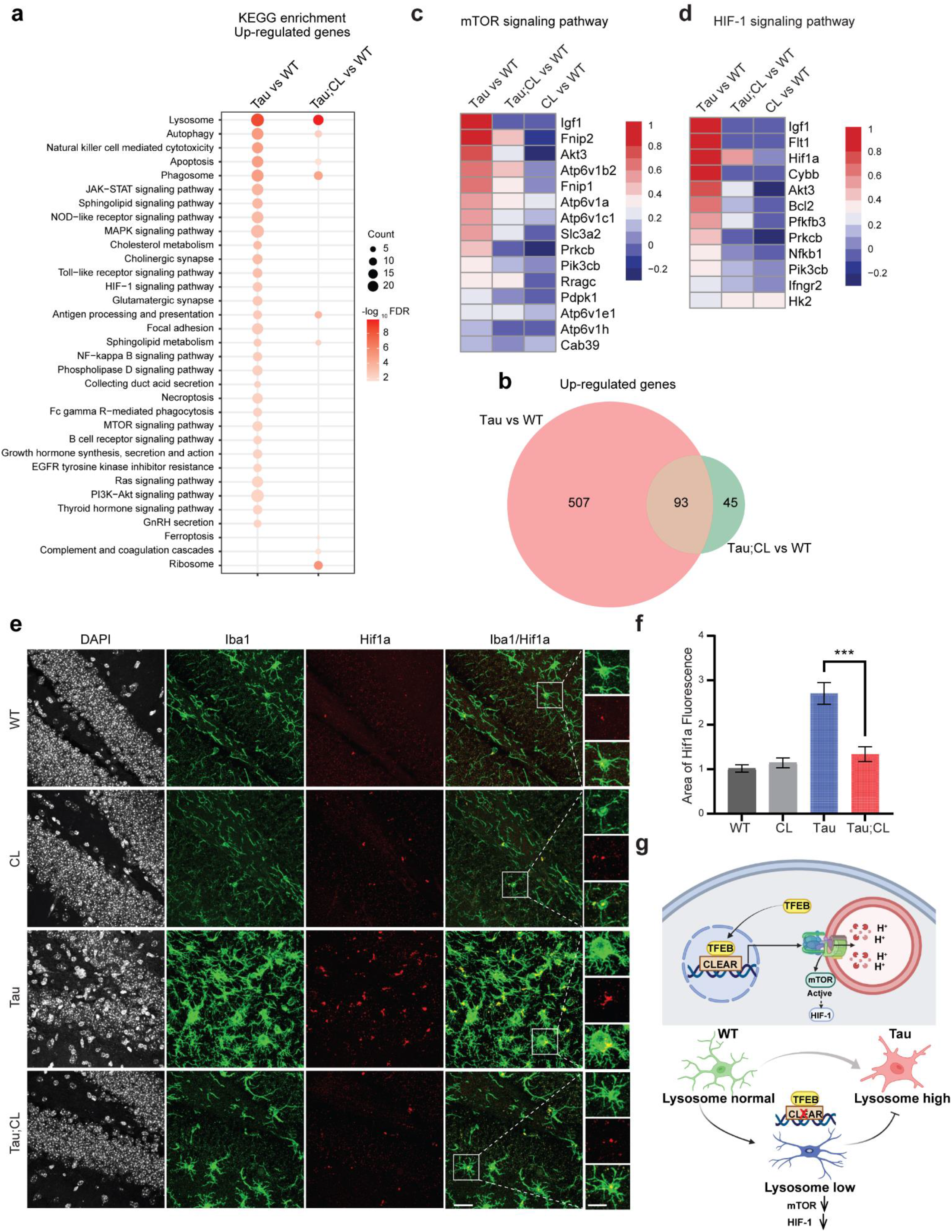
Disruption of TFEB-v-ATPase regulation leads to compromised lysosomal and inflammatory changes and reduced mTOR and HIF-1 signaling in tauopathy. **a**. KEGG enrichment analysis for up-regulated genes comparing Tau;CL with Tau. **b**. Venn diagram summarizing the numbers of up-regulated genes for all microglia in Tau and Tau;CL. **c**,**d**. Heatmaps comparing mTOR (c) and HIF-1 (d) signaling pathway related genes (log_2_ fold change) between different genotypes. **e**. Representative images of 9-month-old WT, CL, Tau and Tau;CL brains stained for HIF1α (red) and Iba-1 (green). Scale bar 50 µm and 25 µm in brackets. **f**. Quantification of percent area fluorescence of HIF1α staining in WT, CL, Tau and Tau;CL mice (N=4). **g**. Diagram depicting the mechanism of microglia activation in Tau mice mediated by TFEB-v-ATPase lysosomal regulation. Data are presented as average ± SEM. Two-tailed Student’s *t-*test. ***p<0.001. See also Extended Data Fig. 7.

Taken together, our data demonstrate that impairment of TFEB regulation of lysosomal acidification and function through v-ATPase transcriptional activation locks microglia in the resting state with downregulated mTOR and HIF-1 metabolic pathways, revealing an essential role of TFEB-v-ATPase lysosomal regulation in microglia function, particularly in activation of innate immune response under stress conditions triggered by tau perturbation, resulting in overall defective microglial response to tau pathology (Fig. 8g).

## Discussion

The lysosome plays essential roles in cellular metabolism and clearance through coordinated lysosome to nucleus signaling. TFEB is a critical regulator in this process, through which it mediates the degradation of protein aggregates characteristic of AD and other neurodegenerative diseases^24^. However, besides the lysosomal genes, TFEB also regulates the transcription of a broad range of other targets^25, 26^, making it difficult to decipher a lysosome-specific mechanism. To tackle this problem, we chose to specifically manipulate TFEB-v-ATPase signaling, given the crucial role of the v-ATPase in regulating lysosomal pH and function. Here we report that the TFEB-v-ATPase transcriptional program is essential in maintaining lysosomal homeostasis under physiological conditions and is required to induce microglial activation in tauopathy. Disruption of the signaling pathway leads to impaired lysosomal activity, heightened tau pathology and failed microglial response to the pathogenic insult.

We demonstrate that TFEB binds to the CLEAR sequence in the *Atp6v1h* promoter and mediates its transcription. This represents part of its expression regulation as mutagenizing the CLEAR sequence only results in ∼30% reduction of *Atp6v1h* mRNA levels. Remarkably, this reduction is sufficient to alter the entire v-ATPase activity and lysosomal pH, highlighting an obligatory role of the V1H subunit in v-ATPase assembly and a physiological function of TFEB in v-ATPase and associated lysosomal regulation. Since many other v-ATPase subunit genes are also TFEB targets, it is possible that manipulating the CLEAR sequence of other v-ATPase targets will have similar effects but this remains to be tested. It is important to note that it is TFEB-V1H/v-ATPase signaling, but not mere gene expression, which confers this effect as a 50% reduction in *Atp6v1h* mRNA in the germline heterozygous mice (VKO) did not display any phenotypes, likely due to genetic compensation. Combined with the *Drosophila* study^14, 15^, our results enforce the notion that TFEB-v-ATPase transcriptional regulation represents an evolutionarily conserved signaling pathway in maintaining lysosomal homeostasis.

Our bulk brain RNA-seq analysis identified upregulation of TFEB and the lysosomal pathway in Tau mice. Using in vitro assays, we showed that insoluble tau promotes TFEB nuclear translocation and downstream gene expression. This effect may be caused directly by the intracellular tau aggregates as TFEB can be activated by various cellular stress and damage signals^25^. Alternatively, this may be triggered by tau induced lysosomal stress as tau is known to be degraded in the lysosome^11^. Although the precise mechanism remains to be established, the fact that reducing the v-ATPase and lysosomal pathway in CL leads to worsened tau pathology supports the idea that the lysosomal pathway upregulation in Tau mice represents an innate adaptive response against pathological tau accumulation. However, this protective mechanism may no longer be effective under chronic tauopathy conditions, necessitating the addition of exogenous TFEB to resolve the increasing tau burden^8, 12^.

Further analysis by snRNA-seq revealed drastically altered microglia profiles in Tau mice, with diminished homeostatic microglia (subcluster 0) and corresponding expansion of transitional subcluster 1, which then converts to two distinct subpopulations: DAM/MGnD-like (subcluster 2) and Interferon-responsive (subcluster 3). These are associated with robust upregulation of lysosomal pathway genes, along with immune and inflammatory genes. The elevated lysosomal pathway is also a prominent feature of DAM ^17^, suggesting that it is a component of the general microglia activation program in response to pathological stimuli in the brain. Critically, our finding that tuning down the lysosomal pathway in Tau;CL microglia leads to impaired glial and immune activation demonstrates that lysosomal pathway upregulation is required to induce microglia activation. In this regard, TFEB has been shown to influence immune response through its regulation of the autophagy-lysosomal pathway and by direct transcriptional activation of immune target genes^26^. Although these mechanisms may indeed be at play, our data that immune activity can be directly regulated by specific TFEB-v-ATPase lysosomal signaling without affecting TFEB or its inflammation targets highlight an essential role of the lysosome in immune system regulation.

Subclustering analysis of WT and CL microglia allowed us to further divide the homeostatic subcluster into 2 populations, 0a and 0b, with 0b enriched in CL microglia. This subcluster expresses low lysosomal genes and is associated with reduced mTOR and HIF-1 signaling pathways. Significantly, subcluster 0b failed to be converted to activated states on Tau background and, therefore, is locked in the homeostatic state. Indeed, the microglial phenotypes observed in the Tau;CL mice recapitulate key features of Trem2 knockout on amyloid mouse models, with both displaying defective microglia activation, reduced DAM signatures and impaired mTOR activity^27, 28^. Since we did not detect changes of Trem2 expression in Tau microglia, we propose that mTOR may serve as a common mediator converging membrane receptor signaling and lysosomal activity to microglial activation.

mTOR plays a central role in cellular metabolism in multiple cell types, including innate immune cells, through regulating several downstream pathways, among them HIF-1 signaling^29, 30^. The activation of microglia requires a metabolic switch from oxidative phosphorylation to aerobic glycolysis to swiftly generate energy for fulfillment of energy-intensive processes such as migration, cytokine production and secretion, phagocytosis and proliferation. Hif1a is a canonical modulator of the metabolic reprogramming^31^. We found significantly increased HIF-1 signaling pathway genes in fully activated microglia subclusters and elevated Hif1a immunostaining in microglia of Tau mice, both of which were reduced by CL, supporting an involvement of the HIF-1 pathway in microglial activation. As a mTOR downstream target, reduced HIF-1 signaling may be caused by dampened mTOR although it is also possible that this event is mTOR independent. Regardless, the markedly reduced mTOR pathway in the microglia subcluster enriched in CL and Tau;CL mice indicate that TFEB-v-ATPase dysregulation not only affects lysosomal acidification and degradative capacity but also impairs mTOR activation, resulting in its inability to undergo the metabolomic reprogramming required for microglia activation.

It is well-established that TFEB responds to mTOR and the LYNUS machinery composed of v-ATPase to undergo cytoplasmic to nucleus trafficking. Our work reveals that nuclear TFEB regulates the v-ATPase transcriptional program, which in turn feedback to regulate lysosomal pH and function. This pathway is not only important for intraneuronal tau clearance but also required for microglia activation in response to tau pathology. These findings demonstrate a critical role of the lysosome, in part modulated by TFEB-v-ATPase signaling, in both neuronal and immune cell function in physiology and diseases of tauopathy.

## Methods

### Animals

All protocols involving mice were approved by the Institutional Animal Care and Use Committee of Baylor College of Medicine. PS19 (Tau) mice were obtained from Jackson Labs ^32^. Heterozygotes were bred to B6C3F1/J wild type mice to maintain the line. CL mice were generated utilizing CRISPR-mediated mutagenesis as described below. Mice were backcrossed to C57BL/6 mice for a minimum of 10 generations.

### CRISPR/Cas9-mediated mutagenesis design and CL mouse production

To introduce the mutagenized CLEAR sequence site in *Atp6v1h*, a single guide RNA (sgRNAs) was selected using the Wellcome Trust Sanger Institute Genome Editing website (http://www.sanger.ac.uk/htgt/wge/), so that a double strand break by the resulting sgRNA/Cas9 complex would be created as proximal to the CLEAR sequence site as possible (https://www.sanger.ac.uk/htgt/wge/crispr/300195847). Homology-mediated repair of the double strand break would be directed by a single-stranded donor DNA containing the mutagenized CLEAR site. The sgRNA was synthesized using DNA templates for in vitro transcription. DNA templates were produced using overlapping oligonucleotides in a high-fidelity PCR reaction ^33^. The PCR products were first purified using the QiaQuick PCR purification kit and used as a template for in vitro transcription of the sgRNA with the MEGAshort script T7 kit (ThermoFisher, AM1354). Following in vitro transcription, RNA was purified using the MEGAclear Transcription Clean-Up Kit (ThermoFisher AM1908). All samples were analyzed by Nanodrop to determine concentration and visualized using the Qiaxcel Advanced System using the RNA QC V2.0 kit to check the quality of RNA product before storage at -80°C. A custom Ultramer® DNA oligonucleotide was purchased from Integrated DNA Technologies (Coralville, IA). Cas9 mRNA was purchased from ThermoFisher (A25640). The sgRNA was reanalyzed by Nanodrop prior to assembling the microinjection mixtures, which consisted of Cas9 mRNA (100ng/μL), sgRNA (20 ng/μL, each), and the donor DNA (100 ng/µL) in a final volume of 60 μL 1xPBS (RNAse-free).

C57BL/6N female mice at 24 to 32 days old were injected with 5 IU/mouse of pregnant mare serum, followed 46.5 hr later with 5 IU/mouse of human chorionic gonadotropin. The females were then mated to C57BL/6J males. Fertilized oocytes were collected at 0.5 dpc for microinjection. The BCM Genetically Engineered Mouse Core microinjected the sgRNA/Cas9/ssOligo mixture into the cytoplasm of at least 200 pronuclear stage zygotes. Injected zygotes were transferred into pseudopregnant ICR females on the afternoon of the injection, approximately 25-32 zygotes per recipient female.

To determine if the mutagenized CLEAR site had been introduced by HDR, N0 mice were genotyped by standard PCR. Two primers approximately 100-200 bases outside the CLEAR site were designed to amplify an amplicon for direct Sanger sequencing. Sequence traces were compared to wild-type DNA to confirm incorporation of the modified bases.

The *Atp6v1h* germline heterozygous mouse was produced by the Baylor College of Medicine Knockout Mouse Phenotyping Program (KOMP2) (https://commonfund.nih.gov/KOMP2). Specifically, exon 3, representing a critical region of the *Atp6v1h* gene, was deleted by employing two Cas9-RGN guides, one each targeting the flanking introns to this critical region. The mice were produced as described above.

The primers for mouse genotyping is listed in Supplementary Table 3.

### Bulk RNA-seq and analysis

RNA was isolated from hippocampal tissues of 4- and 9-month-old WT and Tau mice using RNeasy Mini kit from Qiagen with DNase digestion. cDNA library was generated using the QuantSeq 3′ mRNA-Seq Library Prep Kit following the manufacturer’s instructions. Briefly, oligo(dT) beads were used to enrich mRNA. After chemical fragmentation, the cDNA libraries were generated using NEBNext Ultra RNA Library Prep Kit for Illumina (New England Biolabs) and were assessed using Qubit 2.0 fluorometer to calculate the concentrations and Bioanalyzer Instrument to determine insert size. cDNA library samples were then sequenced using Illumina HiSeq2000 machine with a depth of 50-55 million pairs of reads per sample (Sequencing and Microarray Facility, MD Anderson, Houston, TX). bcl2fastq was used for demultiplexing.

Cutadapt ^34^ was used to remove adapters and low-quality reads. Then, remaining reads were mapped to the mm10 genome using STAR ^35^. Only unique mapped reads were kept for further analysis. Gene counts were produced using featureCounts with default parameters, except for ‘stranded’ which was set to ‘0’. The DESeq2 package was used to identify differentially expressed genes with the cutoff: | log_2_(fold change) | >= 0.5 and FDR<0.05 (Tau (9-month-old) versus WT (9-month-old) or Tau (4-month-old) versus WT (4-month-old)).

Gene Set Enrichment analyses of GO and KEGG were performed by using GSEA v4.3.2 for all expressed genes between Tau group and WT group. Enrichment pathways were ranked based on normalized enrichment score (NES). GO enrichment analysis was done by using Metascape (http://metascape.org/) online tool with default parameters.

### snRNA-seq and analysis

9-month-old WT, CL, Tau and Tau;CL mice were perfused transcardially with cold saline under anesthesia. Hippocampal tissues were dissected into 1.5ml RNAase free Eppendorf tube, flash-frozen with liquid nitrogen, and stored at –80 °C. Single-nucleus suspensions were prepared as described ^36^.

Nuclei stained by Hoechst-33342 were collected using the SONY SH800 FACS sorter. For each 10x Genomics run, 100k–400k nuclei were collected. 10k nuclei for each channel were loaded to the 10x controller. snRNA-seq was performed using the 10x Genomics system with 3’ v3.1 kits. All PCR reactions were performed using the Biorad C1000 Touch Thermal cycler with 96-Deep Well Reaction Module. 13 cycles were used for cDNA amplification and 16 cycles were used for sample index PCR. As per 10x protocol, 1:10 dilutions of amplified cDNA and final libraries were evaluated on a bioanalyzer. Each library was diluted to 4 nM, and equal volumes of 18 libraries were pooled for each NovaSeq S4 sequencing run. Pools were sequenced using 100 cycle run kits and the Single Index configuration. Read 1, Index 1 (i7), and Read 2 are 28 bp, 8 bp and 91 bp respectively. A PhiX control library was spiked in at 0.2 to 1% concentration. Libraries were sequenced on the NovaSeq 6000 Sequencing System (Illumina).

Raw reads demultiplexed by bcl2fastq were mapped to the mm10 genome using CellRanger v.6.0.1 with default parameters. Quality control filtering, variable gene selection, dimensionality reduction, and clustering for cells were conducted using the Seurat v.4.0.6 package. To filter low-quality cells, we removed cells for which less than 200 genes were detected or cells that contained greater than 10% of genes from the mitochondrial genome. Genes expressed in fewer than 3 cells were filtered out. DoubletFinder v.2.0 was used to remove Doublets. Batch effect was corrected by Harmony. Gene expression count data for all samples was normalized with “NormalizedData” function, following by scaling to regress UMIs by “ScaleData” function. Principal component analysis (PCA) and UMAP implemented in the “RunPCA” and “RunUMAP” functions were used to identify the deviations among cells, respectively. For subtypes differential expression markers or genes were identified by using the Wilcoxon test implemented in the “FindMarkers” function, which was considered significant with an average fold change of at least 0.25 and Padj <0.05.

### qPCR and ChIP-qPCR

For qPCR, total RNA was extracted from cell culture using a RNeasy Mini Kit (Qiagen) and cDNA was synthesized from 500 ng total RNA using SuperScript III First-Strand Synthesis System (Invitrogen). For hippocampal brain samples, TRIzol reagent (Invitrogen) was used to extract total RNA and cDNA was synthesized from 2µg total RNA. cDNA was diluted to 2 ng/µL and 4 µL were added to 10 µL 2x FastStart Universal SYBR Green PCR Master (Roche). Each sample was run in triplicate using iTaq Universal SYBR Green Supermix (BioRad, #172-5124) on a CFX384 Touch Real-Time PCR Detection System. Ct values were normalized to the housekeeping gene GAPDH, which was amplified in parallel. The 2^-ΔΔCT method was utilized to calculate relative gene expression levels. The primer sequences are shown in Supplementary Table 4.

For CHIP-qPCR, N2a cells were plated in 10% FBS DMEM and allowed to grow for 48 hours prior to transfection with TFEB-3XFLAG or GFP plasmids. Cells were transfected according to the manufacturer’s protocol at a µL lipofectamine: µg plasmid ratio of 3:1 (X-tremeGENE 9, Roche). After 48 hours, chromatin was isolated (Active Motif high sensitivity ChIP kit) and sheared (Diagenode Bioruptor bath sonicator) using 20 cycles (30 seconds on, 30 seconds off). Chromatin immunoprecipitation was performed using a mouse anti-FLAG antibody (Sigma) or normal mouse IgG (Millipore). Immunoprecipitated DNA was amplified using qPCR primer sets shown in Supplementary Table 5. Data are reported as fold-change of TFEB binding normalized to input and IgG control immunoprecipitation.

### Luciferase assay

HEK293 cells or N2a cells grown in 12-well plates were co-transfected with TFEB-3XFLAG expression vector and Atp6v1h wild type promoter or CLEAR mutant promoter firefly luciferase plasmid, together with Renilla-TK luciferase vector using X-tremeGENE 9 transfection reagent (Roche). The Atp6v1h promoters were cloned into pGL3 plasmid (Promega). The CLEAR mutant promoters were generated using site-directed mutagenesis (QuikChange II XL Site-Directed Mutagenesis Kit, Agilent). Renilla plasmid was transfected at 1/20 the amount of the other plasmids. In studying the effect of tau on Atp6v1h promoter, HEK293 cells were co-transfected with the firefly luciferase Atp6v1h promoter construct, the TauP301L-V5 plasmid, and the Renilla-TK construct. After 24 hours, Pff were added to seed insoluble tau. At 48 hours, cells were lysed in passive lysis buffer (Promega). The Dual-Glo Luciferase Assay System (Promega) was used to determine firefly and Renilla luciferase activities according to the manufacturer’s instructions. Measurements were performed in a white 96 well-plate on a Tecan Spark 10M plate reader.

### In vitro tau seeding assay

Procedure was described in detail in a previous study ^16^. HEK293 cells were grown in DMEM (Life Technologies) with 10% FBS at 37 °C with 5% CO_2_. Cells were cultured in 60 mm^2^ dish with 5 mL medium. At 60% confluency, cells were transfected with Tau P301L-V5 encoding full length human tau with P301L mutation and V5 tag (GKPIPNPLLGLDST) and TFEB-GFP at a 2:1 ratio of tau:TFEB (X-tremeGENE 9, Roche). 24 hours later the media was changed and 40 µL of Pff was added to the culture along with 200 nM Bafilomycin (Sigma) or an equivalent volume of DMSO. 24-48 hours after seeding, cells were collected for analysis.

### Primary mixed glia and microglia cultures and treatment

Primary glia cultures were prepared as described previously ^37^. Briefly, the cerebral cortices were isolated from P3 newborn pups in ice-cold dissection medium [Hanks’ balanced salt solution (HBSS) with 10 mM HEPES, 0.6% glucose, and 1% (v/v) penicillin/streptomycin], with meninges removed. The tissue was then finely minced and digested in 0.125% trypsin at 37°C for 15 min, followed by the addition of trypsin inhibitor (40 μg/mL) and DNase (250 μg/mL). Next, tissue was triturated, and resuspended in DMEM with 10% FBS. The cell suspension was centrifuged and resuspended one more time to remove tissue debris. Cells were plated 24 well culture plates with poly-D-lysine (PDL) coated cover slip at a density of 50,000 cells/cm^2^ and cultured in DMEM with 10% FBS at 37°C in a humidified atmosphere of 95% air and 5% CO2 for 7-10 days. For microglia cultures, suspended cells were plated on T-75 flasks at a density of 50,000 cells/cm^2^ to generate mixed glial cultures. After the mixed glial culture reached confluency, the flasks were shaken for 2 hours at 250 rpm at 37°C. The T-75 flasks were then tapped vigorously 10-15 times on the bench top to loosen microglia growing on top of the astrocytes. The media along with floating microglia were collected from the flask and centrifuged for 5 minutes at 1000 x g. The cell pellet from one T-75 flask was resuspended and plated in PDL coated 24 well plates. The media was changed 24 hours later. After 48-72 hours in culture, microglia were collected for experiments.

For amino acid and serum starvation, cells were incubated in pre-warmed EBSS (Earls Balanced Salt Solution) (Invitrogen) at 37°C for 4 hours to induce autophagy. LysoSensor Green DND-189 stock solution (ThermoFisher) was diluted to the final working concentration (1 μM) in either normal cell culture medium or EBSS. The cells were stained with 1 μM LysoSensor in media for 5 min. Cells were rinsed twice with 1X PBS and incubated in culture medium for confocal microscopy. For LPS treatment, 200 ng/ml LPS (Sigma-Aldrich) was added to the microglia culture media 16 h before an experiment. Overall lysosomal hydrolytic activity was determined with DQ(tm)-BSA dye (Invitrogen). DQ(tm)-BSA stock solutions were prepared according to the manufacturer’s instructions. Cells were incubated with DQ-BSA dye (10 μg/mL) for 16 h. Images were taken using a Confocal microscope and the fluorescence intensities of DQ-BSA were quantified with Fiji (ImageJ).

Acidic Nanoparticles (NP) were prepared as described ^38^. Briefly, Resomer® RG 503H PLGA (Sigma-Aldrich, 719870) was used with a lactide-glycoside ratio of 50:50, to prepare a stock solution of polymer with fluorophore by dissolving 10 mg of PLGA and 0.3 mg of Nile red fluorophore (Sigma-Aldrich, 19123) in 1 mL of tetrahydrofuran (THF, Sigma-Aldrich, 401757). The working solution was made by diluting 100 μL of the stock solution into 10 mL of deionized water under sonication. PLGA-aNP solutions were used as freshly prepared for all experiments and added to culture medium for 16 hours.

### Immunoblotting

For Western blot, cells, forebrain, or dissected hippocampus were lysed in RIPA buffer (TBS with 1% NP-40, 1% sodium deoxycholic acid, 0.1% sodium dodecyl sulfate, and protease/phosphatase inhibitor cocktails (Roche)). Lysates were sonicated 6 pulses at 50% duty cycle and incubated on ice for 30 minutes. Samples were then centrifuged at 20,000 x g for 20 minutes. Supernatants were collected and quantified using a Pierce BCA Protein Assay Kit (Thermo Fisher). Lysates were incubated for 7 minutes at 90°C in sample loading buffer. Fifteen microgram protein samples were loaded onto 12% SDS-PAGE gels, then transferred to nitrocellulose membranes (Bio-Rad). Membranes were blocked in 5% nonfat milk in PBS + 0.1% Tween 20 (PBS-T). Blots were probed with primary antibody, washed with PBS-T, then probed with the appropriate HRP-conjugated secondary antibody, followed by additional washes. The signal was developed with Pierce ECL Western Blotting Substrate (Thermo Fisher). Band intensity was quantified using ImageJ software (National Institute of Health) and normalized to the loading control (β-tubulin).

### Immunostaining

Primary cultures grown on coverslips were fixed in 4% paraformaldehyde (PFA) for 20 minutes at room temperature after multiple washes with ice cold PBS. Following fixation, coverslips were gently washed with PBS. Coverslips were then incubated in blocking buffer (PBS + 2% donkey serum + 0.1% Triton X-100) for one hour at room temperature. After blocking, coverslips were incubated with primary antibodies overnight in blocking buffer at 4°C. Coverslips were then washed in PBS followed by incubation with secondary antibodies for 2 hours in blocking buffer at room temperature. Coverslips were then washed in PBS and mounted using DAPI containing mounting media. Cells were imaged by confocal microscopy (Leica TCS SPE).

Animals were perfused transcardially with 4% PFA in 0.1 M PBS, pH 7.4, under ketamine (300 mg/kg) and xylazine (30 mg/kg) anesthesia. Brains were harvested, post-fixed in the same fixative overnight at 4 °C, dehydrated with 30% sucrose in PBS, and serially sectioned at 30 μm on a sliding microtome (Leica). For immunofluorescence, sections were permeabilized in PBS/0.1% Triton X-100 for 30 min and blocked with 4% normal donkey serum in PBS/0.1% Triton X-100 for 1 h at room temperature. Sections were then incubated with primary antibodies in 2% serum in PBS/0.1% Triton X-100 overnight at 4 °C. Sections were then washed and incubated with Alexa Fluor 488-or Alexa Fluor 555-conjugated secondary antibodies (Invitrogen) for 1 h at room temperature. After washing with PBS, sections were incubated with DAPI to stain the nucleus. Images were captured using a Laser-Scanning Confocal Microscopy (Leica) and quantified with ImageJ.

### Immunofluorescence quantification

TFEB nuclear localization was calculated based on counting instances of DAPI and FLAG staining colocalization and divided by total number of FLAG positive cells per confocal image. Colocalization was determined based on multiple Z-stack slices (20 slices per 30 µm section).

For calculating area fluorescence of AT8, GFAP, and Iba1 antibody staining, the slide containing representative slices of the entire mouse brain was scanned on an EVOS fluorescence microscope. Area fluorescence in specific brain regions was calculated after thresholding to eliminate background and nonspecific staining using ImageJ. Area fluorescence of AT8, GFAP, or Iba1 staining in the hippocampus was averaged across all consistently represented sections for each animal to signify the relative pathology or gliosis within the entire volume of the brain region analyzed.

For microglia morphology quantification, Iba1 positive microglia were imaged by confocal microscopy using a 63x oil lens to generate Z-stacks of the tissue thickness (∼30 μm) with a step-size of 0.5 μm. Z-stacks were analyzed using IMARIS software, in which the Filament function was used to generate filaments for individual cells in the images and microglia processes were automatically rendered based on the Iba1 signal.

### Antibodies

MC1, CP13 and PHF1antibodies were generous gifts from the late Peter Davies (Albert Einstein College of Medicine). All other antibodies used for immunoblotting and staining were purchased from commercial sources described in Supplementary Table 6.

### Statistics

The statistical methods used for bulk and single nuclear RNA-seq are described in their perspective sections. For others, data are presented as average ± standard error of the mean (S.E.M.). Power analysis was performed using a confidence interval of α=0.05. Violin plots are presented as medians and quartiles. Pairwise comparisons were analyzed using a two-tailed Student’s *t-*test. Grouped comparisons were made by one way ANOVA with Sidak’s correction. P-values less than 0.05 were considered statistically significant (*p<0.05, **p<0.01, ***p<0.001).

## Data availability

Bulk hippocampus RNA-seq and snRNA-seq data generated in this study have been deposited in GEO with accession number: GSE218728 (https://www.ncbi.nlm.nih.gov/geo/query/acc.cgi?acc=GSE218728; reviewer token: sxolugighdyltij). Data will be made publicly available as of the date of publication. Any additional information on sequencing data reported in this paper is available upon request.

## Acknowledgements

We are grateful to the Baylor College of Medicine Knockout Mouse Phenotyping Program (KOMP2) and the Genetically Engineered Rodent Models Core for the creation of CL and VKO mice and Cytometry and Cell Sorting Core for FACS analysis. We thank A. Cole, B. Reeves and B. Contreras for expert technical support and members of the Zheng laboratory for stimulating discussions. HL is a CPRIT Scholar in Cancer Research (RR200063). This study was supported by grants from the NIH (P01 AG066606, RF1 NS093652, RF1 AG020670 and RF1 AG062257 to HZ and R00 AG062746 to HL).

## Author contributions

BW, HMS and HZ conceived the project; MS and HL provided input and expertise in CL mutagenesis and snRNA-seq respectively. HMS performed bulk brain RNA-seq, created CL mice and was responsible for initial set of cell and mouse experiments. BW carried out follow up molecular, cellular and biochemical analyses and worked with CQ, ZL, YQ and HL in the snRNA-seq experiments and data analysis. SW assisted in mouse breeding and biochemical analysis, WX constructed acidic nanoparticles and YX performed the seeding experiment. BW, HMS and CQ prepared the figures and BW and HZ wrote the manuscript. All authors read, edited and approved the final manuscript.

