## Supplemental Figures 1-7 for "TFEB-vacuolar ATPase signaling regulates lysosomal function and microglial activation in tauopathy"

### Extended Data Figures and Figure Legends

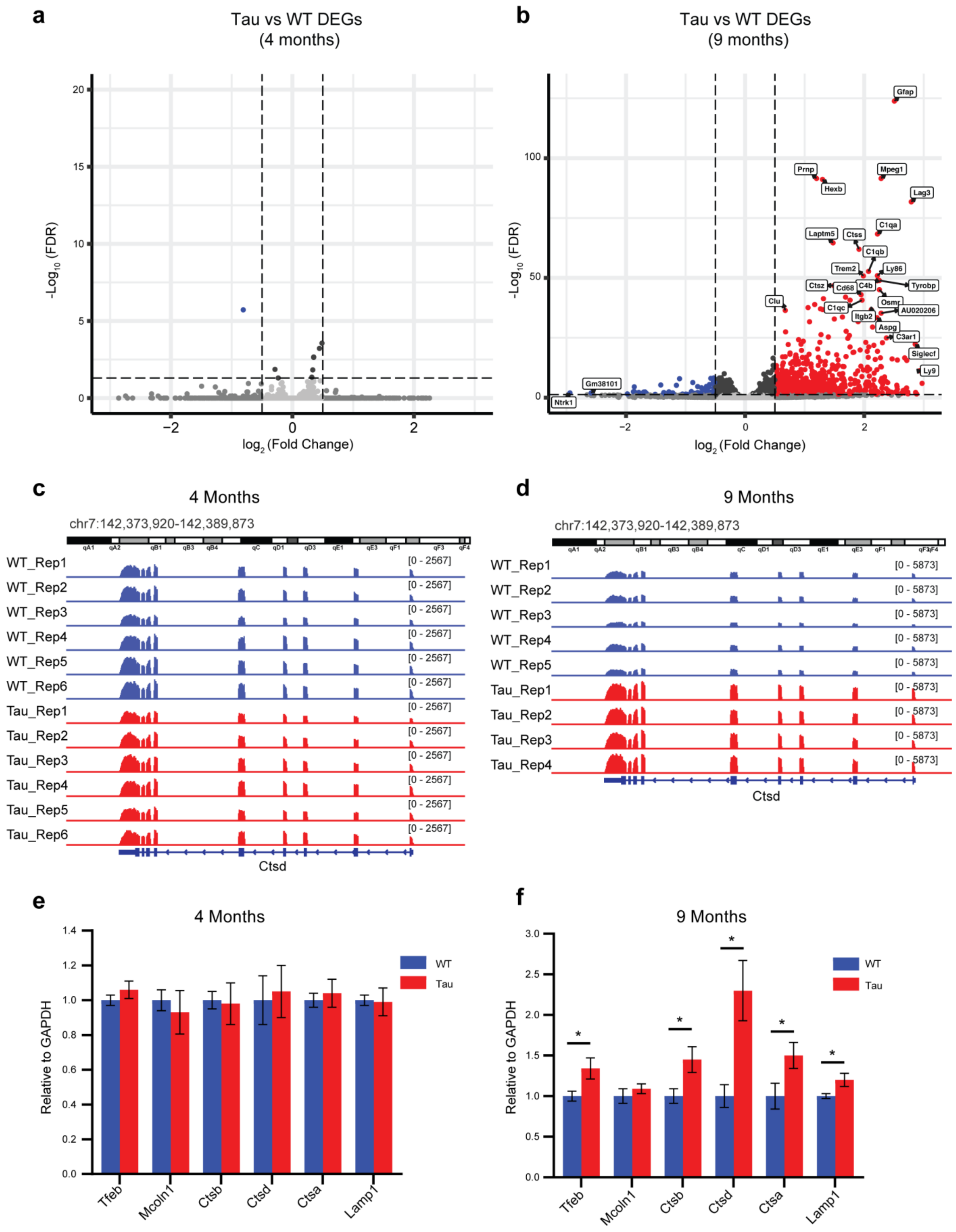

a
Tau vs WT DEGs
(4 months)
-Log10 (FDR)
log2 (Fold Change)
b
Tau vs WT DEGs
(9 months)
Gfap
Prnp
Mpeg1
Hexb
Lag3
C1qa
Laptm5
Ctss
C1qb
Trem2
Ly86
Ctsz
Cd68
C4b
Tyrobp
Clu
Osmr
C1qc
AU020206
Itgb2
Aspg
C3ar1
Siglecf
Ly9
Gm38101
Ntrk1
-Log10 (FDR)
log2 (Fold Change)
c
4 Months
chr7:142,373,920-142,389,873
WT_Rep1
WT_Rep2
WT_Rep3
WT_Rep4
WT_Rep5
WT_Rep6
Tau_Rep1
Tau_Rep2
Tau_Rep3
Tau_Rep4
Tau_Rep5
Tau_Rep6
[0 - 2567]
Ctsd
d
9 Months
chr7:142,373,920-142,389,873
WT_Rep1
WT_Rep2
WT_Rep3
WT_Rep4
WT_Rep5
Tau_Rep1
Tau_Rep2
Tau_Rep3
Tau_Rep4
[0 - 5873]
Ctsd
e
4 Months
Relative to GAPDH
WT
Tau
Tfeb
Mcoln1
Ctsb
Ctsd
Ctsa
Lamp1
f
9 Months
Relative to GAPDH
WT
Tau
Tfeb
Mcoln1
Ctsb
Ctsd
Ctsa
Lamp1

**Figure 1. Lysosomal and immune pathway genes are enriched in 9-month-old Tau mice.** **a**,**b**. Volcano plots showing differentially expressed genes (DEGs) in Tau mice compared with WT mice at 4 and 9 months of age, respectively. Sample information, raw and unique mapped reads are shown in Supplementary Table 1. **c,d**. Snapshots of genome browser tracks of bulk brain RNA-seq of WT and Tau mice at 4 or 9 months at the *Ctsd* loci. **e,f**. qPCR analysis of hippocampal samples showing TFEB and TFEB regulated lysosomal genes are upregulated in 9 month-old but not 4 month-old Tau mice. Data are presented as average ± SEM. Two-tailed Student's *t*-test. *$p<0.05$. N=5/group.

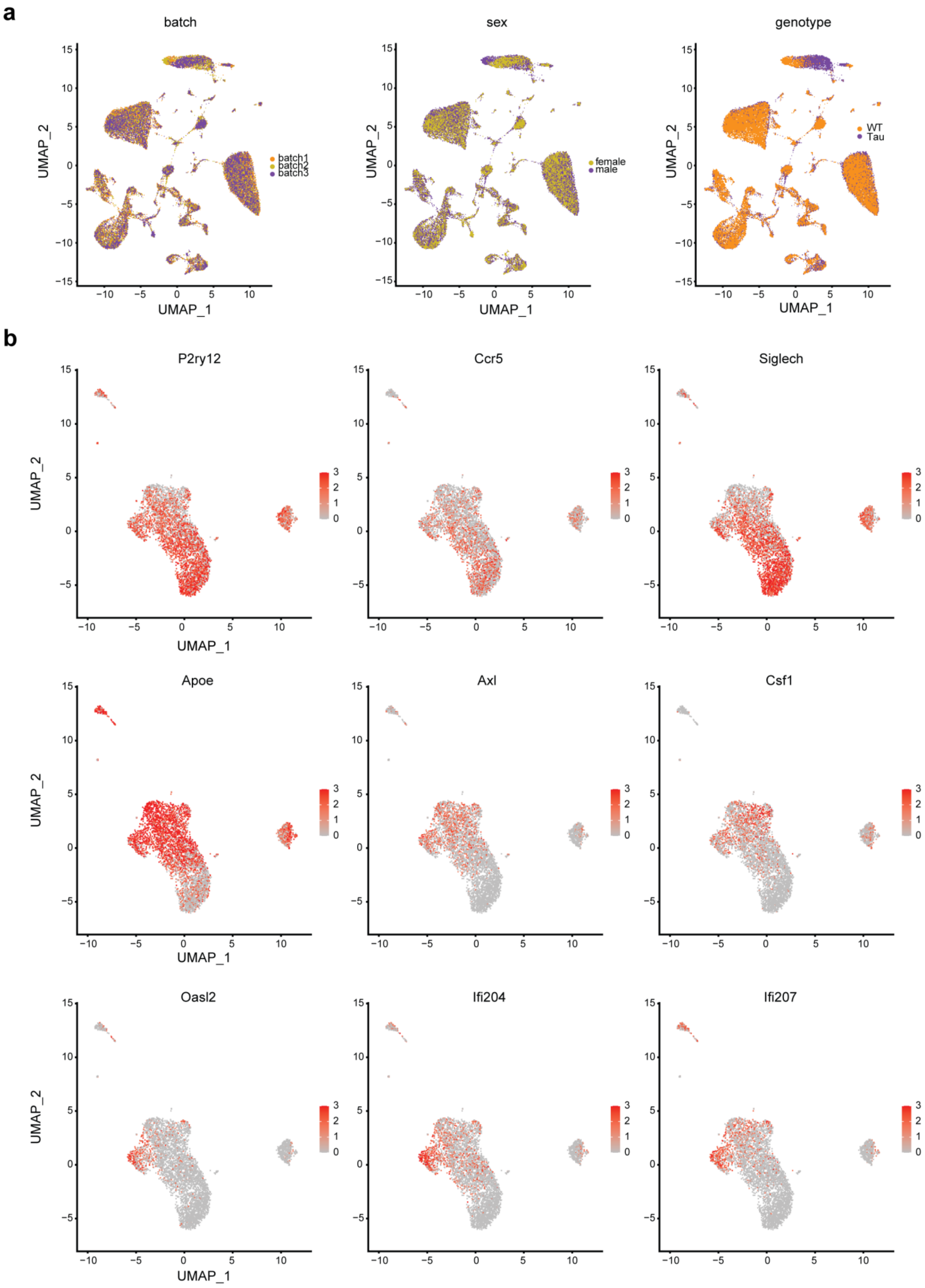
a
batch
sex
genotype
batch1
batch2
batch3
female
male
WT
Tau
UMAP_1
UMAP_2
b
P2ry12
Ccr5
Siglech
Apoe
Axl
Csf1
Oasl2
Ifi204
Ifi207

**Figure 2. snRNAseq characterization of microglia subtypes.**

**a**. UMAP plots of 55,254 cells from hippocampus of WT and Tau mice after batch effect corrections including batch, sex and genotype. **b.** UMAP representation of re-clustered microglia from WT and Tau mice analyzed by snRNA-seq. The expression levels of homeostatic microglia genes (*P2ry12, Ccr5* and *Siglech*), disease-associated-microglia genes (*Apoe, Axl* and *Csf1*) and IFN responsive-microglia genes (*Oasl2, Ifi204* and *Ifi207*) are displayed. Sample information, raw and unique mapped reads are shown in Supplementary Table 2.

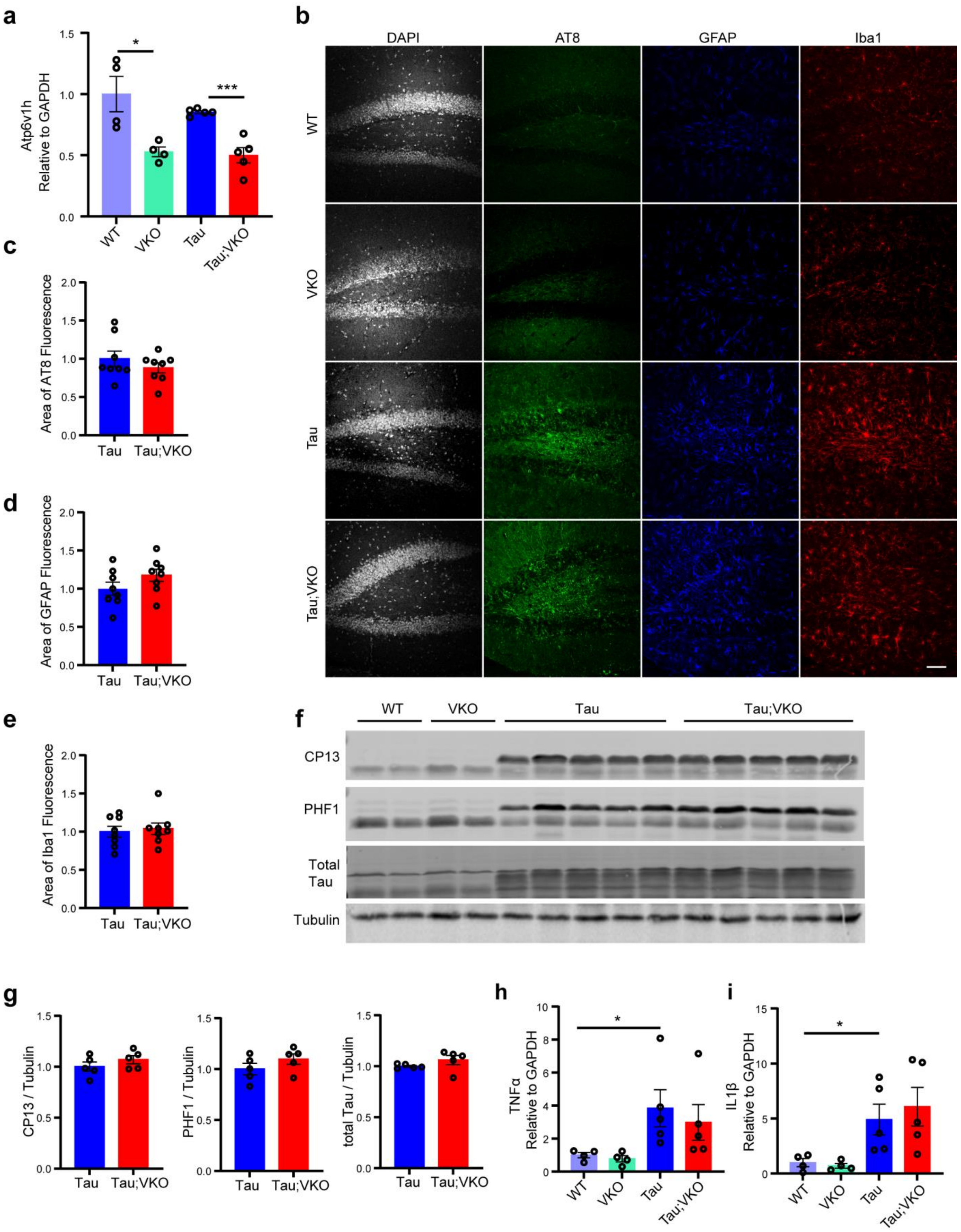

a
b
c
d
e
f
g
h
i
Atp6v1h Relative to GAPDH
WT
VKO
Tau
Tau;VKO
DAPI
AT8
GFAP
Iba1
Area of AT8 Fluorescence
Area of GFAP Fluorescence
Area of Iba1 Fluorescence
CP13
PHF1
Total Tau
Tubulin
CP13 / Tubulin
PHF1 / Tubulin
total Tau / Tubulin
TNFα Relative to GAPDH
IL1β Relative to GAPDH

**Figure 3. *Atp6v1h* heterozygous deletion does not affect phospho-tau levels or gliosis in Tau mice.** **a.** qPCR analysis of *Atp6v1h* transcripts in 9-month-old WT, *Atp6v1h* heterozygous knockout (VKO), Tau and Tau;VKO hippocampus tissues, revealing ~50% reduction of *Atp6v1h* mRNA in VKO and Tau;VKO mice. **b-e**. Representative fluorescent confocal images of AT8, GFAP and Iba1 immunostaining with quantification in the dentate gyrus of 9-month-old WT, VKO, Tau and Tau;VKO. Scale bar: 50 µm. N=8/group. **f,g**. Western blot with quantification of total and phospho-tau species recognized by CP13 and PHF1antibodies. Samples were derived from forebrain lysates of 9-month-old WT, VKO, Tau and Tau;VKO. N=5/group. **h,i**.. qPCR analysis of $TNF\alpha$ and $IL1\beta$ in 9-month-old WT, VKO, Tau and Tau;VKO hippocampus tissues showing no differences between Tau and Tau;VKO samples. N=5/genotype. Data are presented as average ± SEM. Two-tailed Student's *t*-test. *p<0.05, ***p<0.001.

a

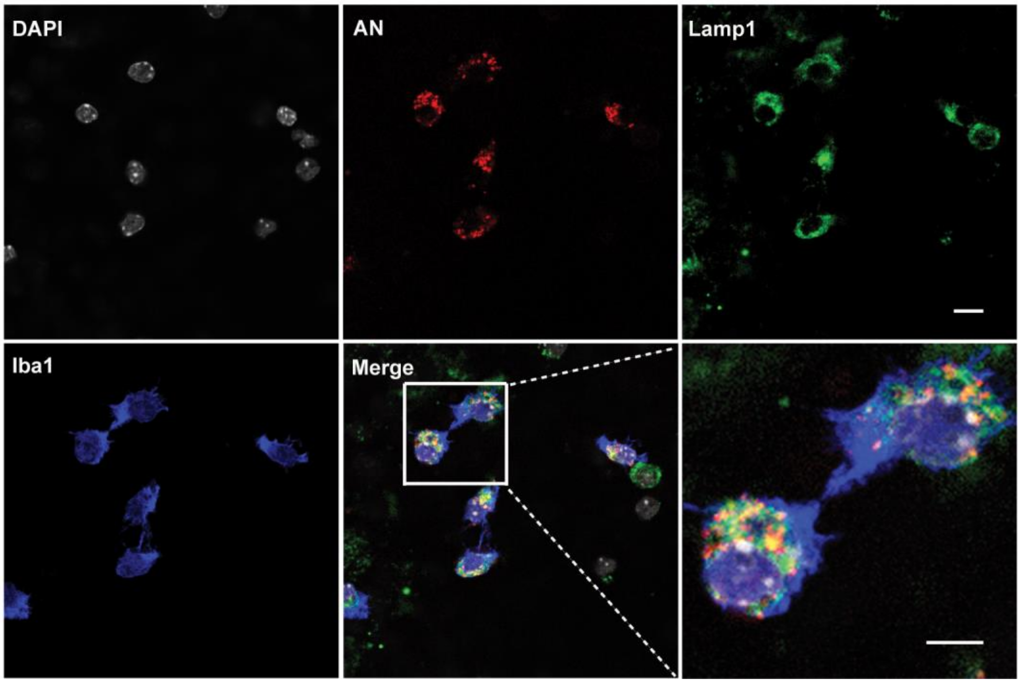


**Figure 4. Acidic nanoparticles are targeted to the lysosomes.** Confocal images showing Acidic nanoparticles (AN) co-localized with Lamp1-marked lysosomes in microglia culture. Scale bar: 10 µm.

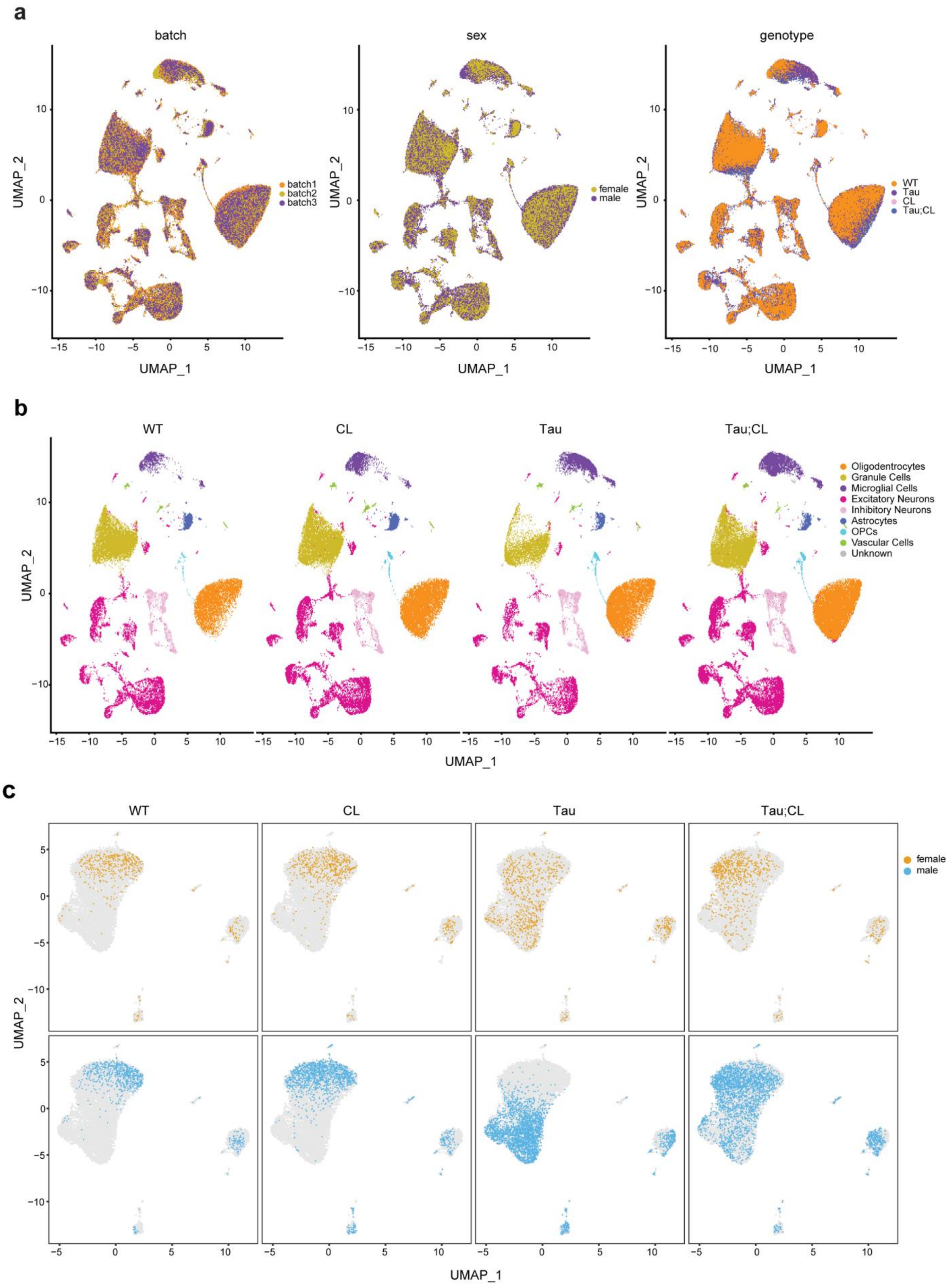
a
batch
sex
genotype
UMAP_2
UMAP_1
batch1
batch2
batch3
female
male
WT
Tau
CL
Tau;CL
b
WT
CL
Tau
Tau;CL
Oligodentrocytes
Granule Cells
Microglial Cells
Excitatory Neurons
Inhibitory Neurons
Astrocytes
OPCs
Vascular Cells
Unknown
UMAP_2
UMAP_1
c
WT
CL
Tau
Tau;CL
female
male
UMAP_2
UMAP_1

**Figure 5. snRNA-seq analysis of hippocampus from WT, Tau, CL and Tau;CL mice.**

**a**. UMAP plots of 137,734 cells from the hippocampus of WT, Tau, CL and Tau;CL mice after batch effect corrections including batch, sex and genotype. **b**. UMAP plots of 137,734 cells from hippocampus across each genotype. **c**. UMAP plots of re-clustered microglia cells in female (orange) and male (blue) across genotypes. Sample information, raw and unique mapped reads are shown in Supplementary Table 2.

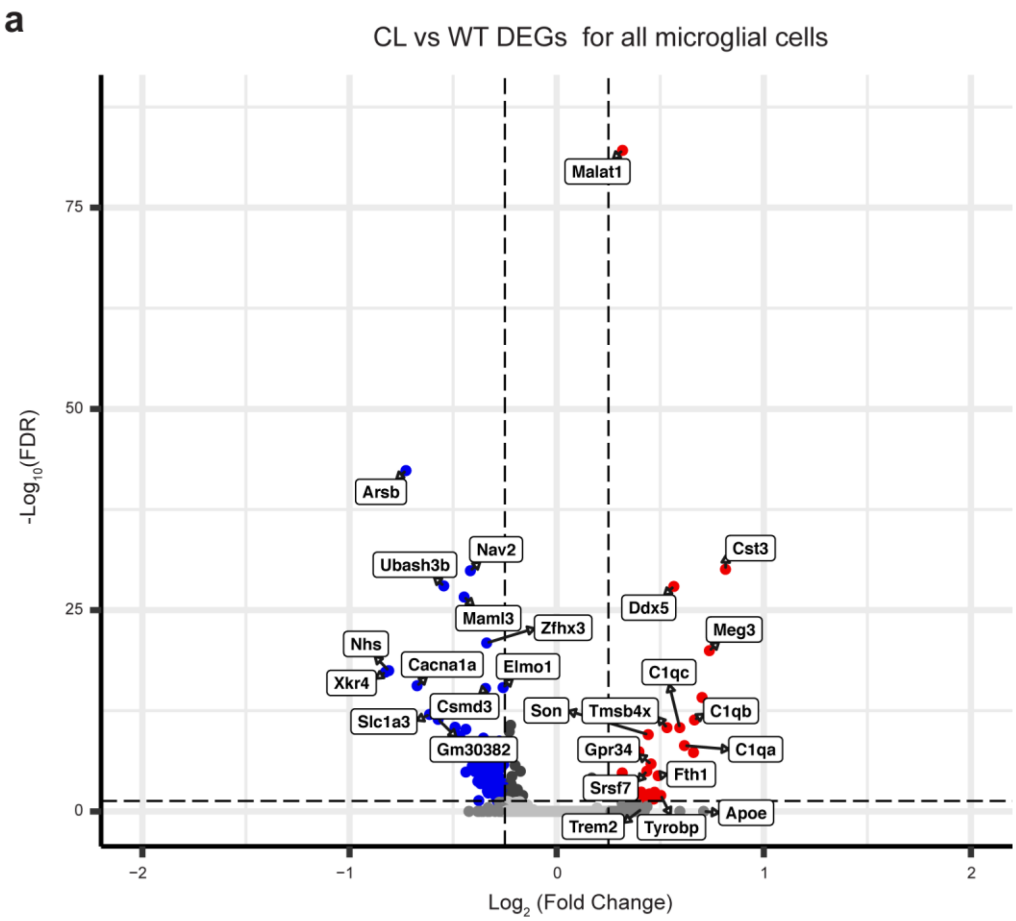


**Figure 6. Differentially expressed genes analysis in CL versus WT microglia.**

Volcano plots showing DEGs for all microglia populations in CL versus WT mice. Up-regulated genes are highlighted in red; Down-regulated genes are highlighted in blue.

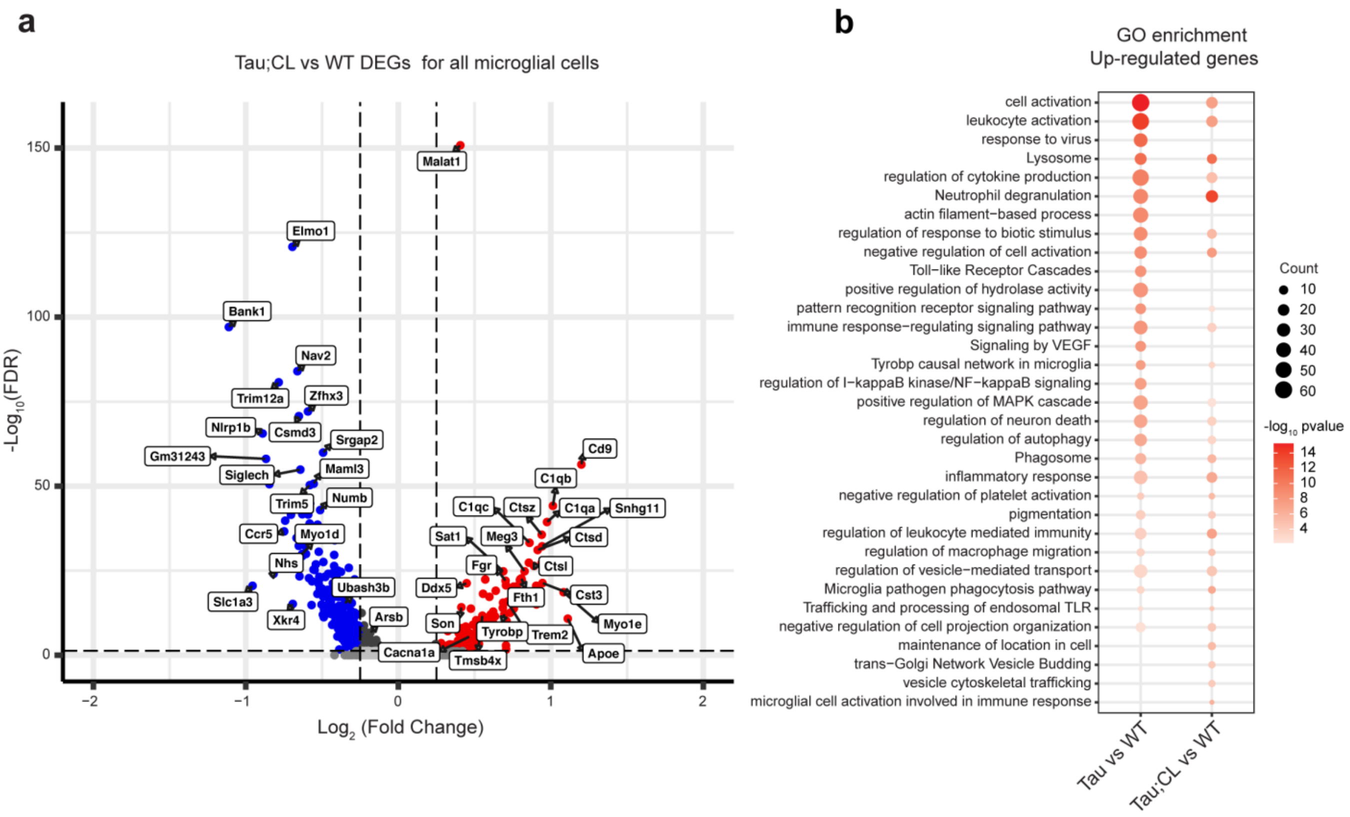


**Figure 7. Reduced lysosomal and inflammatory pathway genes in Tau;CL microglia.**

**a**. Volcano plots showing DEGs for all microglia in Tau;CL versus WT genotype mice. Up-regulated genes are highlighted in red; Down-regulated genes are highlighted in blue. **b.** GO enrichment analysis for up-regulated DEGs in Tau;CL vs WT comparing with Tau vs WT.
